# Thrombospondins 1 and 4 undergo coordinated transport to multicore cytotoxic granules to regulate SMAP biogenesis and function in CTL-mediated cytotoxicity

**DOI:** 10.1101/2024.04.25.590546

**Authors:** Chiara Cassioli, Nagaja Capitani, Claire C. Staton, Claudia Schirra, Francesca Finetti, Anna Onnis, Nadia Alawar, Szu-Min Tu, Ludovica Lopresti, Vanessa Tatangelo, Carmela Tangredi, Salvatore Valvo, Hsin-Fang Chang, Annachiara Miccoli, Ewaldus B. Compeer, Jemma Nicholls, Bruce Blazar, Giuseppe Marotta, Matthew J. A. Wood, Livio Trentin, Laura Patrussi, Michael L. Dustin, Ute Becherer, Cosima T. Baldari

## Abstract

Supramolecular Attack Particles (SMAPs) are particulate entities, characterized by a cytotoxic core enriched in granzymes and perforin surrounded by a glycoproteic shell, released by CTLs and NK cells. Prior proteomic analysis identified thrombospondin-1 (TSP-1) and thrombospondin-4 (TSP-4) as putative components of SMAPs. While TSP-1 has been validated as a component of the SMAP shell and shown to contribute significantly to CTL-mediated killing, the expression and function of TSP-4 in CTLs, and its interplay with TSP-1 in SMAP biogenesis and function, has not been investigated as yet. Here we demonstrate that TSP-4 and TSP-1 have a complementary expression profile during in vitro human CD8^+^ T cell differentiation to CTLs and sequentially localize to lytic granules (LG), with TSP-4 being required for TSP-1 association with LGs. Correlative light microscopy identified the TSP-enriched LGs as the SMAP-containing multicore granules. We show by STED microscopy a heterogeneity among TSP-enriched LGs, the most abundant population being positive for both TSP-4 and TSP-1. We also show that TSP-1 and TSP-4 are co-released in association with SMAPs at immune synapses formed on planar supported lipid bilayers, as assessed by dSTORM imaging. Finally, we provide evidence that TSP-4 is required for CTL- and SMAP-mediated cell killing. Of note, we found that chronic lymphocytic leukemia (CLL) cell supernatants, which suppress CTL mediated killing, also suppress expression of TSP-4 as well as of cytolytic effectors and impair SMAP biogenesis. These results identify TSP-4 as a key player in SMAP structure and activity and suggest that SMAPs may be a new target for immune suppression by CLL.

## Introduction

Cytotoxic T cells (CTLs) eliminate virally infected and tumoral cells using a diversified arsenal of cytotoxic mediators stored in lysosome-related organelles (LRO) that undergo exocytosis upon CTL encounter of cognate cell targets (Cassioli and Baldari, 2022; Chang et al, 2023). Lytic granules (LG) are specialized secretory lysosomes (Peters et al., 1991) that contain perforin (Prf), a pore-forming protein that enables a set of serine proteases, the granzymes (Gzm), to enter the cytoplasm of the target cell and rapidly trigger programmed cell death (Cassioli and Baldari, 2022; McKenzie and Valitutti, 2023). Additionally, CTLs can induce target cell death using membrane-bound FasL (Green and Llambi, 2015), a pool of which is stored in a distinct population of LROs that release it upon CTL activation in association with extracellular vesicles (Bossi et al., 2000). An alternative Prf- and Gzm-dependent mechanism of CTL-mediated cytotoxicity has been recently described, orchestrated by the supramolecular attack particles (SMAP), a class of cytotoxic nanoparticles consisting of a core of Prf and Gzms enclosed in a non-membranous glycoprotein shell (Bálint et al., 2020). The SMAPs are stored in multicore granules (MCG), a class of LGs differing from the canonical single core LGs (SCG) in size, morphology and protein composition (Chang et al., 2022). How SMAPs are targeted to MCGs, how they are assembled and released at the immune synapse (IS) formed by CTLs with their cognate cellular targets, and their specific role versus the know cytotoxic strategies deployed by CTLs, are as yet open questions.

A mass spectrometry-based analysis of SMAPs released by CTLs on functionalized planar supported lipid bilayers (PSLBs), that act as artificial targets that capture released particles, identified thrombospondins (TSP)-1 and -4, but only TSP-1 was subsequently validated as a component of the glycoprotein shell as a ∼60 kDa species that corresponds to its C-terminal portion (Balint et al., 2020). The TSPs are a family of evolutionarily conserved Ca^2+^-binding glycoproteins that, with the exception of specialized cell types such as platelets where they are components of secretory lysosomes (Blair and Flaumenhaft 2009), undergo constitutive secretion to the extracellular matrix (ECM) to regulate cell-cell and cell-matrix interactions (Adams and Lawler, 2011; Henkin and Volpert, 2011; Frangogiannis, 2012; Stenina-Adognravi, 2013). The human TSP family consists of five members that share a C-terminal signature domain, comprising a series of EGF-like domains and Ca^2+^-binding type 3 repeats. The N-terminus of TSPs is more variable and differs in domain organization and degree of oligomerization, which is mediated by a α-helical coiled-coil domain adjacent to the N-terminal laminin binding domain. Group A TSPs are trimeric and include TSP-1 and TSP-2, whereas group B members are pentameric and include TSP-3, TSP-4 and TSP-5 (Lawler et al., 1985; Sottile et al., 1991; Qabar et al., 1995). Additionally, group A TSPs are characterized by central TSP type 1 domains, which are absent in group B TSPs, that inhibit angiogenesis through binding to CD36 and mediate their attachment to multiple cell types (Lawler and Detmar, 2004). TSPs interact with a variety of proteins, including cell surface receptors, ECM components, growth factors and cytokines to form multiprotein complexes that affect a variety of cellular processes (Adams and Lawler, 2011; Henkin and Volpert, 2011; Frangogiannis, 2012; Stenina-Adognravi, 2013). Interestingly, CRISPR/Cas9 targeting of TSP-1 in human CTLs impairs their killing ability (Balint et al., 2020), highlighting a new specific function for this protein in CTL-mediated cytotoxicity.

In addition to TSP-1, another TSP, TSP-4, was identified in SMAPs by mass spectrometry (Balint et al., 2020). The expression and function of TSP-4 in T cells, and its cooperation or redundancy with TSP-1 in SMAP assembly and function, have not been investigated as yet. Here we show that TSP-4 expression is upregulated during CTL differentiation concomitant with a downregulation of TSP-1, localizes to the MCGs with TSP-1 and, similar to TSP-1, participates in CTL-mediated and SMAP-mediated killing. However, TSP-4 is transported to MCGs faster than TSP-1 and is required for TSP-1 localization to MCGs, indicating a distinct and essential role for TSP-4 in SMAP biogenesis. We also provide evidence that SMAPs are targets of soluble factors released by chronic lymphocytic leukemia (CLL) cells to evade CTL-mediated killing, further highlighting SMAPs as key components of the CTL killing arsenal.

## Results

### Complementary profiles of TSP-1 and TSP-4 expression during CD8^+^ T cell differentiation to CTLs

To investigate TSP-4 expression in human CD8^+^ T cells and its regulation during their differentiation to CTLs we first optimized a protocol for in vitro generation of CTLs. Total CD8^+^ T cells purified by negative selection from buffy coats of healthy donors were activated with magnetic beads coated with anti-CD3 and anti-CD28 antibodies in the presence of IL-2 and further expanded for 3 to 5 days after activation (Fig.S1A), when they expressed the typical markers of effector T cells (Fig.S1B) and acquired cytolytic activity as assessed by granzyme B (GzmB) and perforin (Prf) expression (Fig.S1C). Fluorometric-based killing assays performed using MEC1 cells (Stacchini et al., 1999) loaded with a mix of Staphylococcal superantigens (SAg) for polyclonal activation confirmed that these CTLs were able to effectively kill target cells (Fig.S1D).

Expression of TSP-4 (encoded by *THBS4*) during CTL differentiation was compared to that of TSP-1 (encoded by *THBS1*) by RT-qPCR analysis of mRNA purified from total CD8^+^ T cells either freshly isolated (time 0) or at day 3 and 5 after activation (days 5 and 7, see figure S1A). Surprisingly, TSP-1 and TSP-4 expression underwent a differentiation-related regulation in a complementary manner. While detectable at relatively low levels in freshly isolated CD8^+^ T cells, TSP-4 was upregulated during their differentiation to CTLs (Fig.1A). Conversely, TSP-1 was expressed at very high levels in freshly purified total CD8^+^ T cells followed by a substantial relative decrease in CTLs (Fig.1A), although TSP-1 transcripts were still detectable at these stages. This opposite expression pattern was also observed at the protein level by immunoblot. Freshly purified CD8^+^ T cells showed low levels of a ∼150 kDa immunoreactive TSP-4 species that increased in CTLs (Fig.1B,C). Conversely, freshly purified CD8^+^ T cells showed high levels of TSP-1, with a prevalence of the 60 kDa isoform reported to be associated with SMAPs (Balint et al., 2020) that dropped during differentiation to CTLs, although TSP-1 remained detectable at all timepoints analyzed (Fig.1B,C). The specificity of the TSP-1 and TSP-4 immunoreactive bands was confirmed in TSP-1/TSP-4 KO CTLs (Fig.S1E,F). These results demonstrate that TSP-4 is expressed in CTLs, validating the mass spectrometry data (Balint et al., 2020) and, surprisingly, show that expression of TSP-1 and TSP-4 is modulated in opposite directions during CTL differentiation.

**Figure 1.**
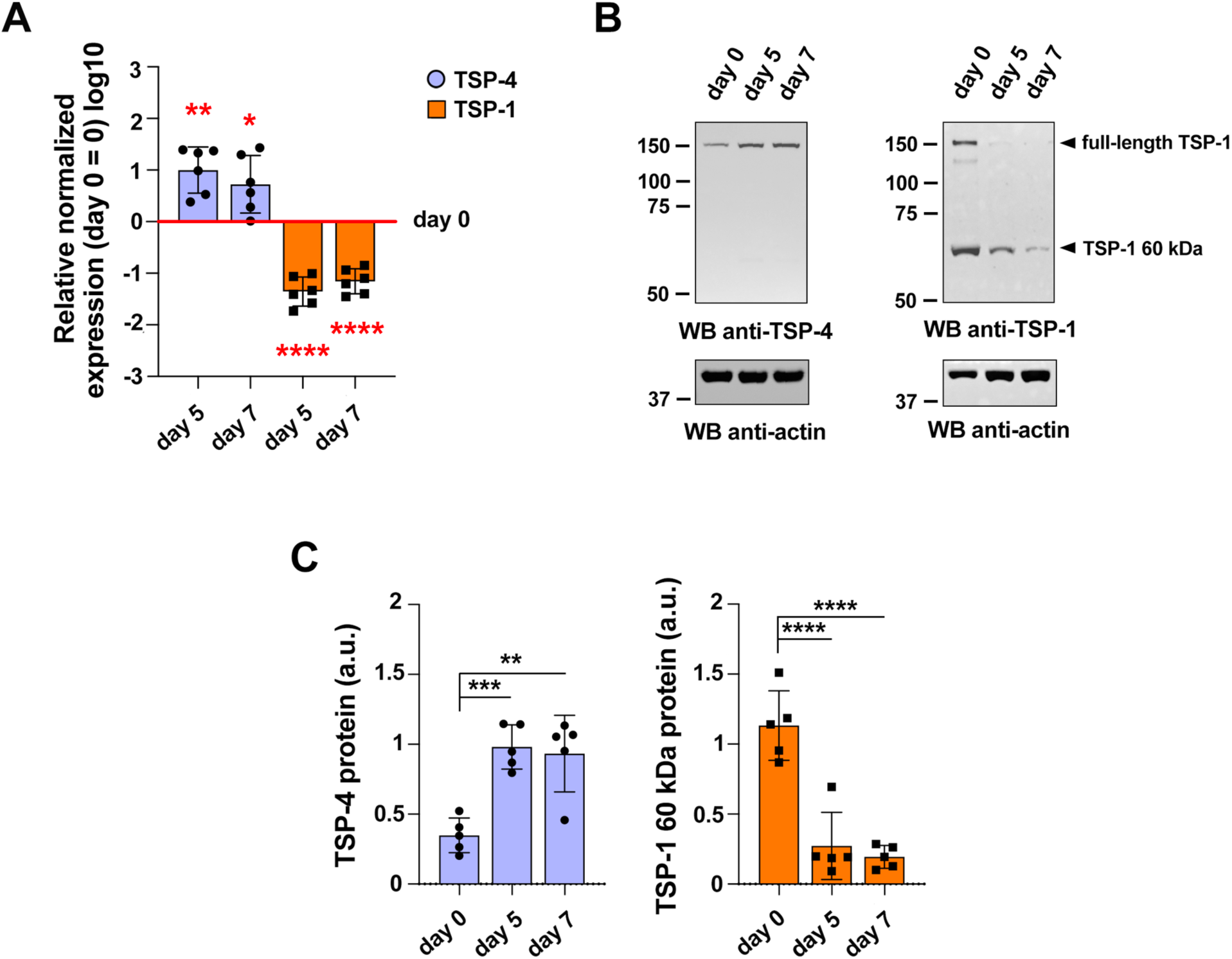
TSP-1 and TSP-4 show an opposite expression profile during CD8^+^ T cell differentiation to CTLs. (**A**) Time course analysis of TSP-4 and TSP-1 mRNA levels in freshly isolated CD8^+^ T cells (day 0) and 5-day and 7-day CTLs. The graph shows the normalized relative abundance (mean±SD, ctr value = 1) of TSP-4 and TSP-1 transcripts. N_donors_ = 6, one-way ANOVA test; **** p ≤ 0.0001, *** p ≤ 0.001, ** p ≤ 0.01, only significant differences are shown. **(B,C**) Immunoblot analysis of TSP-4 (*left*) and TSP-1 (*right*) in lysates of freshly isolated CD8^+^ T cells (day 0) and 5-day and 7-day CTLs. Arrowheads in (*right*) indicate full-length TSP-1 and its 60 kDa species. Quantification (mean±SD) of the relative TSP-4 and 60kDa TSP-1 expression normalized to actin used as loading control is shown in panel C. The migration of molecular mass markers is indicated (kDa). N_donors_ = 5, one-way ANOVA test; **** p ≤ 0.0001, *** p ≤ 0.001, ** p ≤ 0.01, only significant differences are shown.

### TSP-4 co-localizes with TSP-1 in MCGs in CTLs

To analyze the intracellular localization of TSP-4 in CTLs we generated a construct encoding mCherry-tagged TSP-4 (Fig.S2A). Flow cytometric and immunoblot analysis of TSP-4 in CTLs transiently transfected with either the mCherry-encoding vector or the TSP-4-mCherry construct confirmed the expression of recombinant TSP-4-mCherry (Fig.S2B). Additionally, fluorescence microscopy analysis of the CTL transfectants showed a punctate staining for TSP-4-mCherry, as opposed to the expected diffuse cytoplasmic staining of mCherry (Fig.S2F). For comparison, the same analysis was carried out for TSP-1, using GFPSpark-tagged TSP-1 or the respective GFPSpark control (Fig.S2A). The results confirmed the granular pattern of TSP-1-GFPSpark in CTLs (Fig.S2G) (Balint et al., 2020).

To determine the subcellular localization of TSP-4 within the cytolytic endosomal compartment we carried out a confocal immunofluorescence analysis of CTLs transiently transfected with mCherry-tagged TSP-4 and co-stained for the LG markers GzmB, Prf and LAMP-1. The cis-Golgi marker GM130 was used as a negative control. Colocalization analyses showed that TSP-4-mCherry was associated with GzmB, Prf and LAMP-1 positive LGs (Fig.2A). A similar analysis carried out on TSP-1-GFPSpark expressing CTLs (Fig.2B) confirmed the LG localization of TSP-1 (Balint et al, 2020).

**Figure 2.**
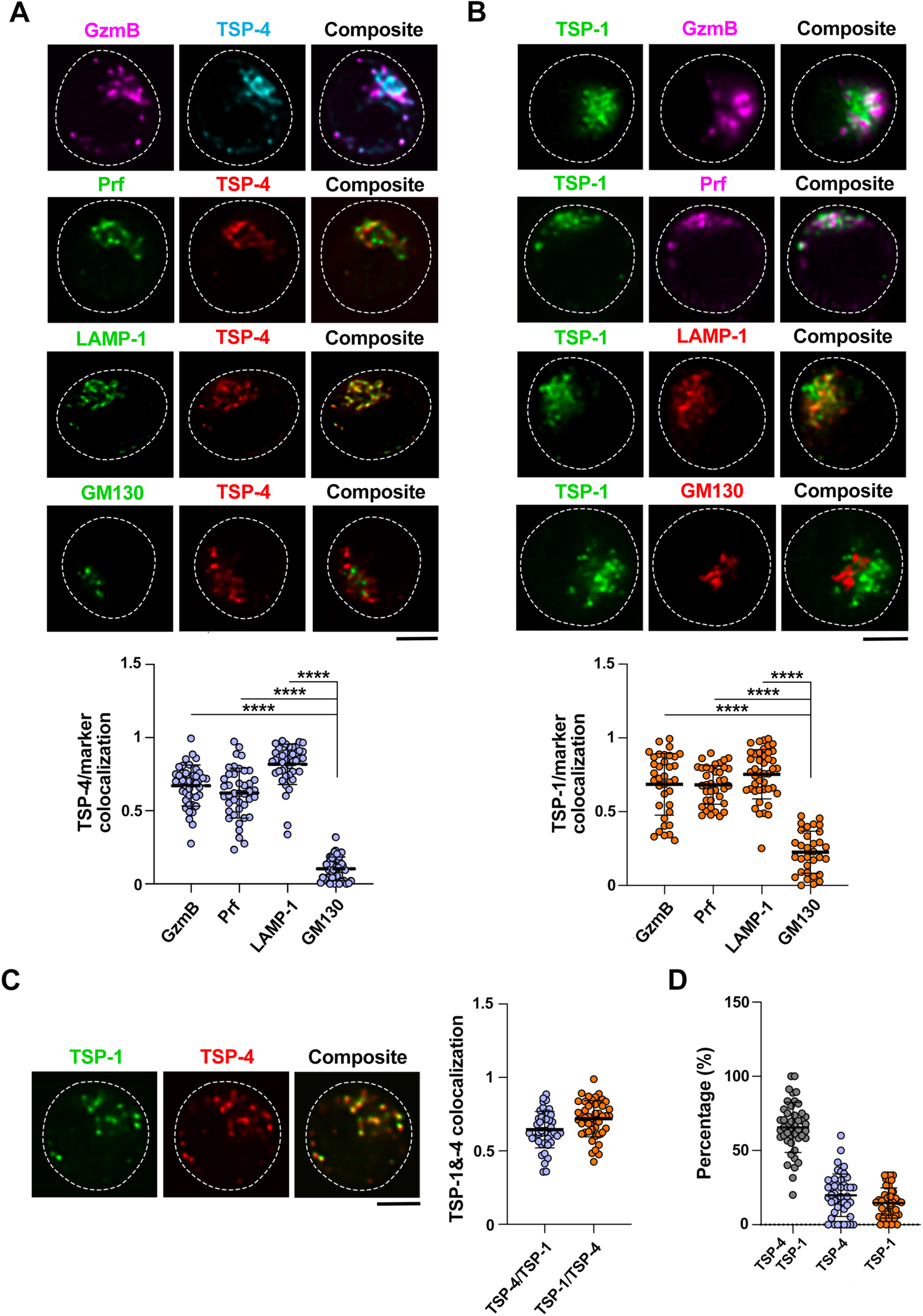
TSP-4 co-localizes with TSP-1 in LGs. (**A,B**) *Top*, Confocal images (medial optical sections) of CTLs expressing either mCherry-tagged TSP-4 (**A**) or GFPSpark-tagged full-length TSP-1 (**B**) co-stained with antibodies against GzmB, Prf, LAMP-1 or GM130. Dashed lines mark the cell outline. Scale bar: 5 μm. *Bottom*, Quantification (mean±SD) of the weighted colocalization using the Manders’ overlap coefficient between TSP-4 (**A**) or TSP-1 (**B**) staining and the signals of each marker. N_donors_ = 3, n_cells_ ≥ 28, Kruskal-Wallis test; **** p ≤ 0.0001, only significant differences are shown. Each dot represents one cell. (**C**) *Left*, Confocal images (medial optical sections) of CTLs transiently co-transfected with constructs encoding TSP-4-mCherry and TSP-1-GFPSpark. Dashed lines mark the cell outline. Scale bar: 5 μm. *Right*, Quantification (mean±SD) of the weighted colocalization using the Manders’ overlap coefficient between TSP-4 and TSP-1 staining N_donors_ = 3, n_cells_ ≥ 28, Kruskal-Wallis test; only significant differences are shown. Each dot represents one cell. (**D**) Quantification (%) of vesicles single or double positive for TSP-4 and TSP-1.

While the mass spectrometry analysis of CTL-derived SMAPs revealed the presence of both TSP-1 and TSP-4 (Balint et al, 2020), whether they co-localize within the same LGs is not known. To address this issue, we co-transfected CTLs with the constructs encoding TSP-4-mCherry and TSP-1-GFPSpark. Transfection efficiency was comparable for the two constructs in these and all other experiments (data not shown). TSP-4 showed a significant vesicular colocalization with TSP-1 (Fig.2C). Accordingly, a substantial subpopulation of LGs contains both TSP-4 and TSP-1 (Fig.2D).

Since TSP-1 is a component of SMAPs (Balint et al., 2020), which are stored in MCGs (Chang et al., 2022), we used TSP-1 to track SMAP biogenesis and follow its interplay with TSP-4 in the MCGs. The vesicular colocalization of TSP-4 with TSP-1 suggests that, similar to TSP-1, TSP-4 may be selectively directed to the MCGs. To test this hypothesis, we performed a Correlative Light and Electron Microscopy (CLEM) analysis of CTLs co-expressing TSP-4-mCherry and TSP-1-GFPSpark. 16 h post-transfection CTLs were incubated for 2 h with fluorescently tagged wheat germ agglutinin (WGA), which labels glycoproteins and has been shown to selectively mark the SMAP-enriched MCGs (Chang et al., 2022). CTLs were then plated on sapphire disc-immobilized anti-CD3 Ab to induce IS formation and LG accumulation. Cells were subjected to high-pressure feezing, freeze-substituted, embedded in resin, sectioned into serial 100 nm sections and subjected to high resolution Structured Illumination microscopy (SIM) and Transmission Electron Microscopy (TEM) analysis (Figure S3A). The fluorescent images were overlaid on the corresponding TEM images to visualize the ultrastructure of the TSP-1, TSP-4 and WGA fluorescent spots. Electron micrographs of human CTLs without electroporation are shown as control (Figure S3B).

The labelled organelles included populations that were either single positive or double positive for TSP-1 and TSP-4 (Fig.3A-C). Within this vesicle pool, our CLEM data demonstrate that the majority of TSP-1 and TSP-4 staining localizes to MCGs (Fig.3B,C), defined as the organelles that encapsulate multiple dense core particles identified as SMAPs (Chang et al., 2022) (Fig.S3B). Consistent with their identity as SMAPs, the average diameter of dense-core particles within the MCGs was 144.26 ± 7.66 nm (Fig.3A,B,E) (Balint et al., 2020; Chang et al., 2022). Interestingly, when fluorescent WGA was fed to live CTLs prior to fixing to mark biosynthetically active MCGs that are still receiving material from the endosomal pathway, these were found to be either TSP-4^+^WGA^+^ or TSP-1^+^TSP-4^+^WGA^+^, while no TSP-1^+^WGA^+^ vesicles could be detected (Fig.3C). This heterogeneity suggests a sequential transport of TSP-4 and TSP-1 to MCGs during their formation, with TSP-1 becoming incorporated at a later step of MCG biogenesis compared to TSP-4. Notably, similar to TSP-1 (Chang et al., 2022), we did not detect TSP-4 associated with single dense cores or compartmentalized within SCGs (Fig.3D), indicating that TSP-4 does not localize to SCGs.

**Figure 3.**
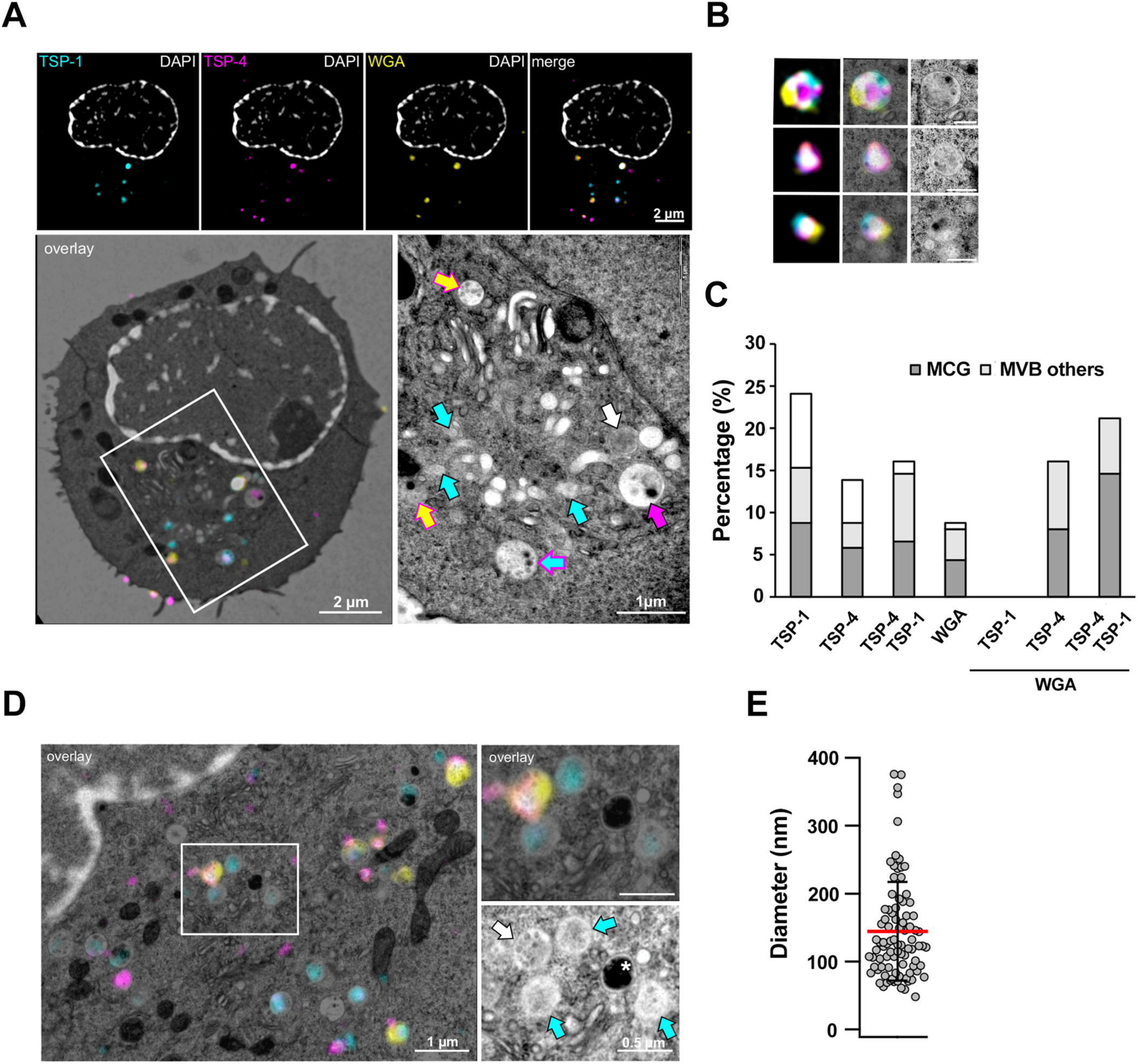
TSP-4, TSP-1 and WGA co-localize in MCGs. (**A**) CLEM image of a representative human CTL expressing TSP-1-GFPSpark and TSP-4-mCherry, preincubated with WGA647 and stained with DAPI. The cell was stimulated by seeding it on an anti-CD3 (UCHT1) coated coverslip. Shown are SIM images (top row) and the corresponding TEM overlay image (bottom panel, left). The white rectangle marks the region, magnified in the lower, right image. Scale bar: 2 µm. In the magnified TEM image (right), arrows point to different fluorescent proteins using the same colour code as the upper panels. Scale bar: 1 µm. (**B**) Enlarged organelles which are positive for TSP-1-GFPspark, TSP-4-mCherry, and WGA647 cropped from the images of three different human CTL. SIM images (left), SIM/TEM overlay images (middle) and TEM images (left). Scale bar: 0.5 µm. (**C**) Quantification of the different organelle fractions in human CTLs analyzed by CLEM, defined by their expression profile and morphology (multicore granule, MCG; multivesicular body, MVB and others). N_donors_ = 2, n_cells_ = 24, 137 organelles. (**D**) Representative CLEM image of a cell section as described in (A) showing a SIM image with the corresponding TEM image (left). Scale bar: 1 µm. The white rectangle marks the region, magnified in the right upper and lower panel. The arrows point to different organelles according to the color code of the fluorescent proteins shown in (A). The white asterisk indicates a SCG. Scale bar: 0.5 µm. (**E**) Quantification of SMAP diameter observed in MCGs as shown in (A and B). N_donors_ = 2, n_cells_ = 24, 90 SMAPs. Red line represents mean and black lines SD.

To analyze the respective arrangement of TSP-4 and TSP-1 within MCGs we exploited the super resolution capability of Stimulated Emission Depletion (STED) microscopy. Day-6 CTLs were co-transfected with constructs encoding TSP-4-HA and TSP-1-FLAG. 16 h post-transfection they were plated on immobilized anti-CD3 Ab and co-stained with antibodies specific for HA, FLAG and GzmB. GzmB was detected in confocal mode, while TSP-1-FLAG and TSP-4-HA were analyzed by STED microscopy. TSP-1 and TSP-4 showed a granular pattern (Fig.4A,B) and co-localized with GzmB in 26.54 ± 2.64 % (n=32) of all granules (Fig.4C). Consistent with the CLEM analysis, the largest population of GzmB^+^ granules (LGs) was TSP-1^+^TSP-4^+^ (52.8 ± 3.2 %, n=32), with a smaller population of TSP-4^+^GzmB^+^ granules and only a minor population of TSP-1^+^GzmB^+^ (Fig.4D), again suggesting an earlier incorporation of TSP-4 into MCGs compared to TSP-1.

**Figure 4.**
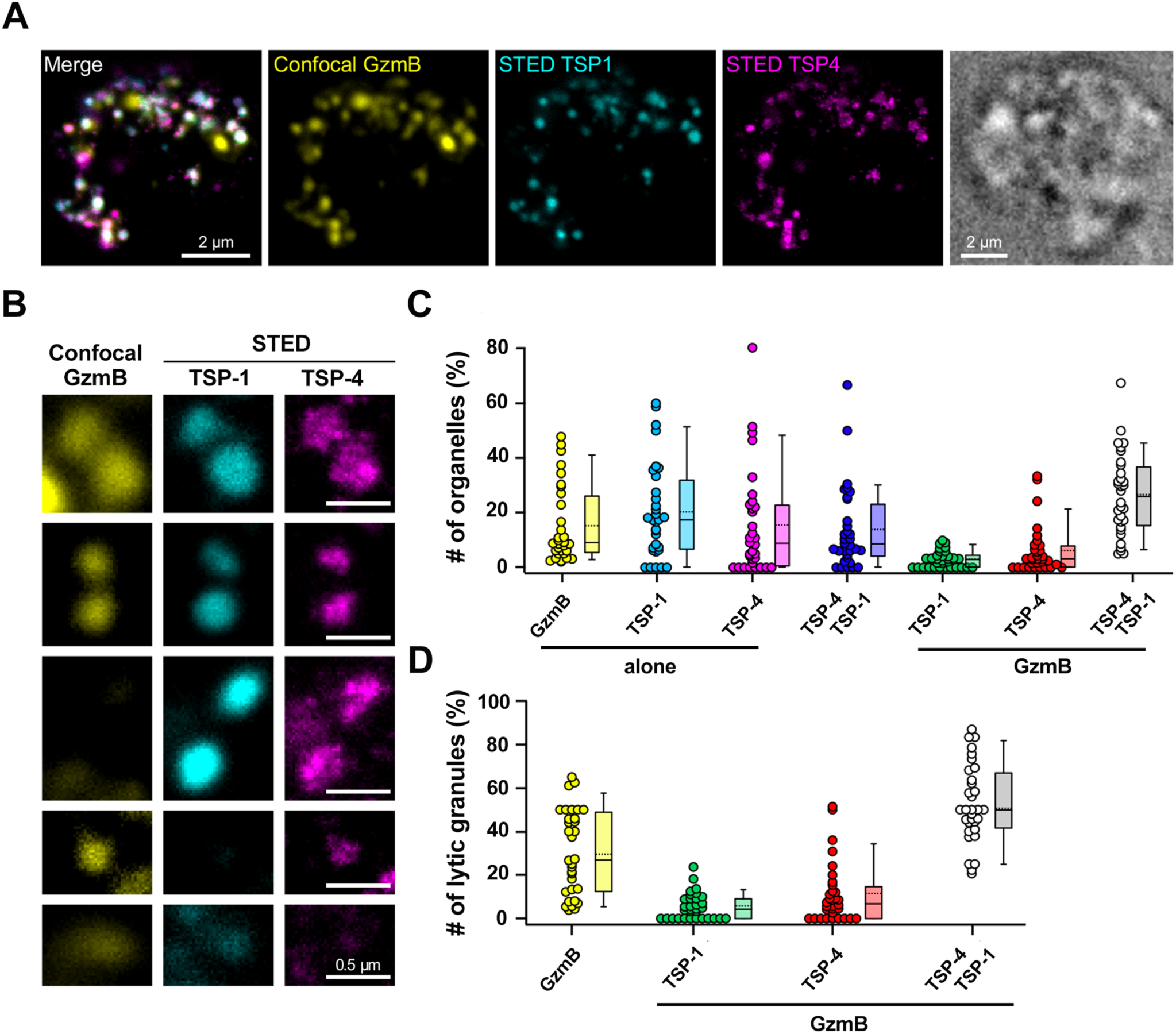
TSP-4 staining has a distinct grainy pattern when localized to LGs. (**A**) Representative stimulated human CTL expressing TSP-1-Flag (cyan) and TSP-4-HA (magenta) forming an IS with the anti-CD3 coated coverslip. Cells were fixed and stained with anti-GzmB (yellow), anti-Flag and anti-HA antibodies. TSP1 and TSP4 were acquired in STED microscopy mode while GzmB was acquired in confocal mode. Shown are the merge image of all three channels and the images of the individual channels. On the right is depicted the brightfield image of the same cell. Scale bar: 2 µm. (**B**) Enlarged pictures of organelles, which are positive for GzmB, TSP-1-Flag and/or TSP-4-HA cut from four different human CTL images as displayed in (**A**). Scale bar: 0.5 µm. (**C,D**) Object-based colocalization analysis of all three proteins displayed as a scatter dot plot with the values of individual cells super imposed with a box plot comprising the median (line) and the mean value (stippled line). While (C) comprises all individual organelles, (D) represents only lytic granules, i.e. organelles that display a GzmB labeling with or without TSP1 and or TSP4. N_donors_ = 2, n_cells_ = 32, 1427 organelles.

### TSP-4 and TSP-1 are sequentially transported to the LGs

To elucidate whether TSP-4 localizes indeed to LGs prior to TSP-1 we used the Retention Using Selective Hooks (RUSH) assay (Boncompain et al., 2012). In this assay the protein of interest is tagged with a fluorescent tag and with streptavidin-binding protein (SBP), which allows its retention in a given membrane compartment through binding to a "hook", an amino acid sequence from a protein resident in that specific compartment, tagged with streptavidin. On addition of biotin to the culture medium, the protein of interest is released from the donor compartment and can be tracked during its transport to its final compartment of destination (Boncompain et al., 2012). We generated two RUSH constructs, encoding either SBP-tagged TSP-1-EGFP or SBP-tagged TSP-4-mCherry, and an endoplasmic reticulum-specific hook (li and KDEL, respectively) (Fig.5A). These constructs were transfected by nucleofection into CTLs and tracked by confocal microscopy to their LAMP-1^+^ LG destination at different time points after addition of biotin. A time course analysis showed an increase in the colocalization of both TSPs with LAMP-1 after biotin addition with TSP-4 arriving prior to TSP-1 (Fig.5B-C), confirming the hypothesis that the two TSPs are sequentially transported to LGs.

**Figure 5.**
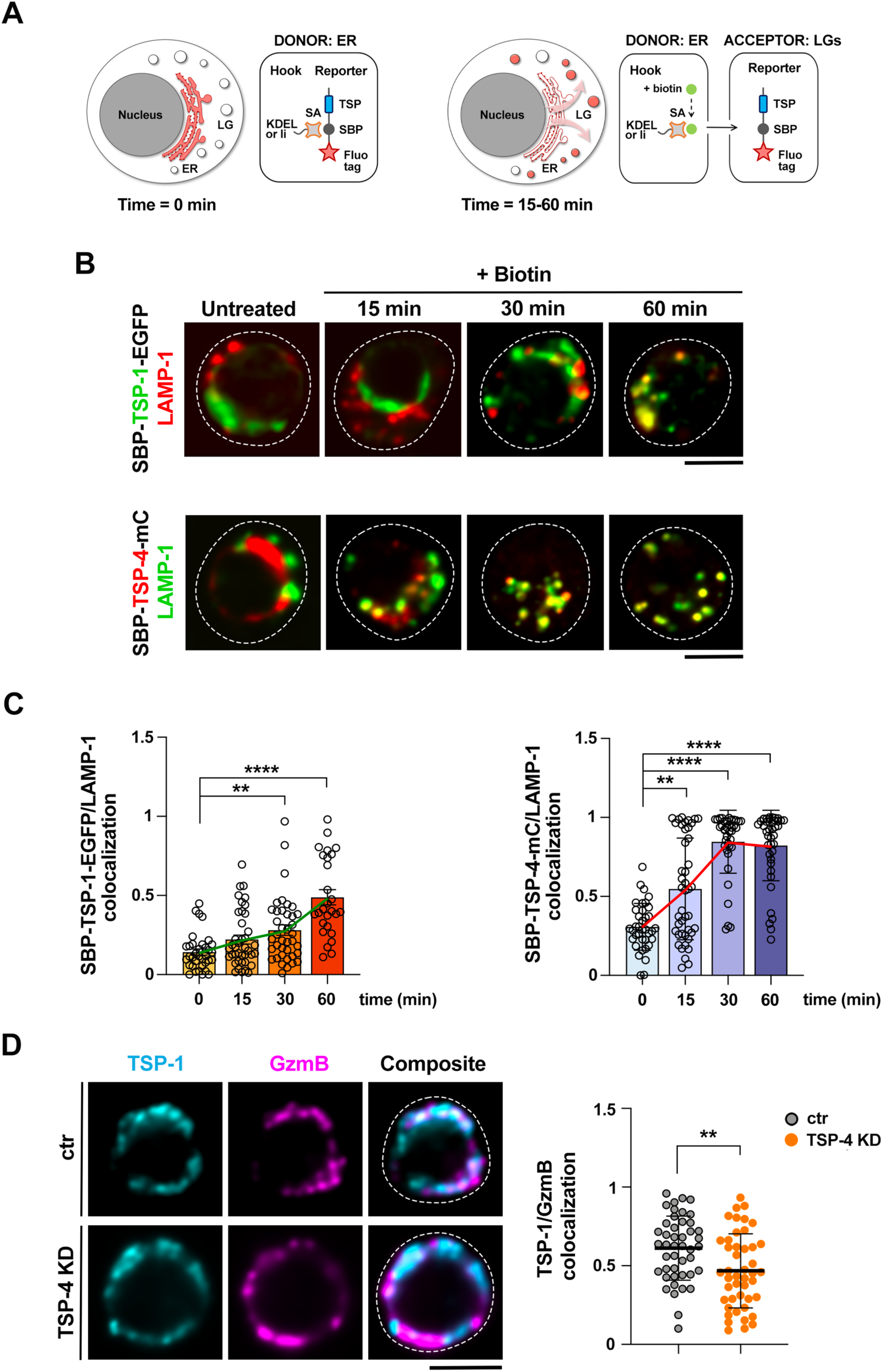
TSP-4 localizes to LGs prior to TSP-1 and is required for TSP-1 association with LGs. (**A**) Principle of the RUSH-secretion assay. In the absence of biotin, TSP-1-streptavidin binding peptide (SBP)-EGFP and TSP-4-SBP-mCherry reporters are retained in the endoplasmic reticulum (ER) by a streptavidin (SA)-li and a streptavidin-KDEL hooks, respectively (left). After biotin addition, TSP-1-SBP-EGFP and TSP-4-SBP-mCherry reporters leave the ER and traffic towards LAMP-1^+^ lytic granules (LGs) (right). (**B**,**C**). Time course analysis in CTLs expressing either Str-KDEL_SBP-TSP-4-mCherry (bottom) or Str-li_SBP-TSP-1-EGFP (top) in the absence or presence of biotin. Representative images (medial optical sections) are shown. Dashed lines mark the cell outline. Scale bar: 5 μm. (**C**) Quantification (mean±SEM) of the weighted colocalization using the Manders’ overlap coefficient between TSP-4 (right) or TSP-1 (left) and LAMP-1 signals. N_donors_ = 2, n_cells_ > 20, Kruskal-Wallis test; **** p ≤ 0.0001, ** p ≤ 0.01, only significant differences are shown. (**D**) Confocal images (medial optical sections) of CTLs expressing TSP-1-GFPSpark and TSP-4-targeting siRNAs (TSP-4 KD) or control scrambled siRNAs. Dashed lines mark the cell outline. Scale bar: 5 μm. Quantification (mean±SD) of the weighted colocalization using the Manders’ overlap coefficient between TSP-1 and GzmB singals. N_donors_ = 3, n_cells_ ≥ 40, Unpaired t test; ** p ≤ 0.01, only significant differences are shown. (**E**) Quantification (%) of vesicles single or double positive for TSP-1GFSpark and GzmB.

The low abundance of TSP-1^+^ LGs compared to TSP-4^+^ and TSP-4^+^TSP-1^+^ ones (Fig.3, Fig.4) raises the possibility that TSP-4 might facilitate the incorporation of TSP-1 into LGs. To test this hypothesis, we investigated the association of TSP-1 with LGs in ctr and TSP-4 KD CTLs that express comparable levels of TSP-1-GFPSpark. The colocalization of TSP-1 with GzmB was significantly decreased in TSP-4 deficient CTLs (Fig.5D,E), indicating that TSP-4 not only localizes to LGs prior to TSP-1, but is required for TSP-1 incorporation into LGs.

### TSP-1 and TSP-4-containing LGs concentrate at the IS

SMAPs are released at the IS formed by CTLs (Balint et al., 2020). To investigate the colocalization of TSP-4 with TSP-1 at the mature IS we plated CTLs transfected with the TSP-1-GFPSpark and TSP-4-mCherry constructs, alone or in combination, on PSLBs functionalized with ICAM-1 to promote LFA-1-mediated adhesion, alone (non-activating conditions) or in combination with an anti-CD3ε Fab’ (activating conditions) (Fig.6A). After 30 min incubation at 37°C, cells were fixed and permeabilized, followed by staining with anti-GzmB antibodies to identify LGs. Consistent with the colocalization analysis of unstimulated CTLs (Fig.2), TSP-4-mCherry showed a significant colocalization with GzmB in single transfectants when plated on non-activating PSLBs (Fig.6B,C), similar to TSP-1-GFPSpark (Fig.6B,C). This colocalization was enhanced under activating conditions, with an accumulation of TSP-4^+^GzmB^+^ vesicles and TSP-1^+^GzmB^+^ vesicles at the IS (Fig.6B,C; videos 1-4). Analysis of double-transfected CTLs confirmed the colocalization of TSP-1 and TSP-4 in LGs that were dispersed within the cell in non-activating conditions and accumulated at the IS in activating conditions (Fig.6B-D; videos 1 and 2). Hence TSP-4 and TSP-1 co-localize under homeostatic conditions as well as during the activation-induced LG maturation events that precede their exocytosis, concentrating at the IS.

**Figure 6.**
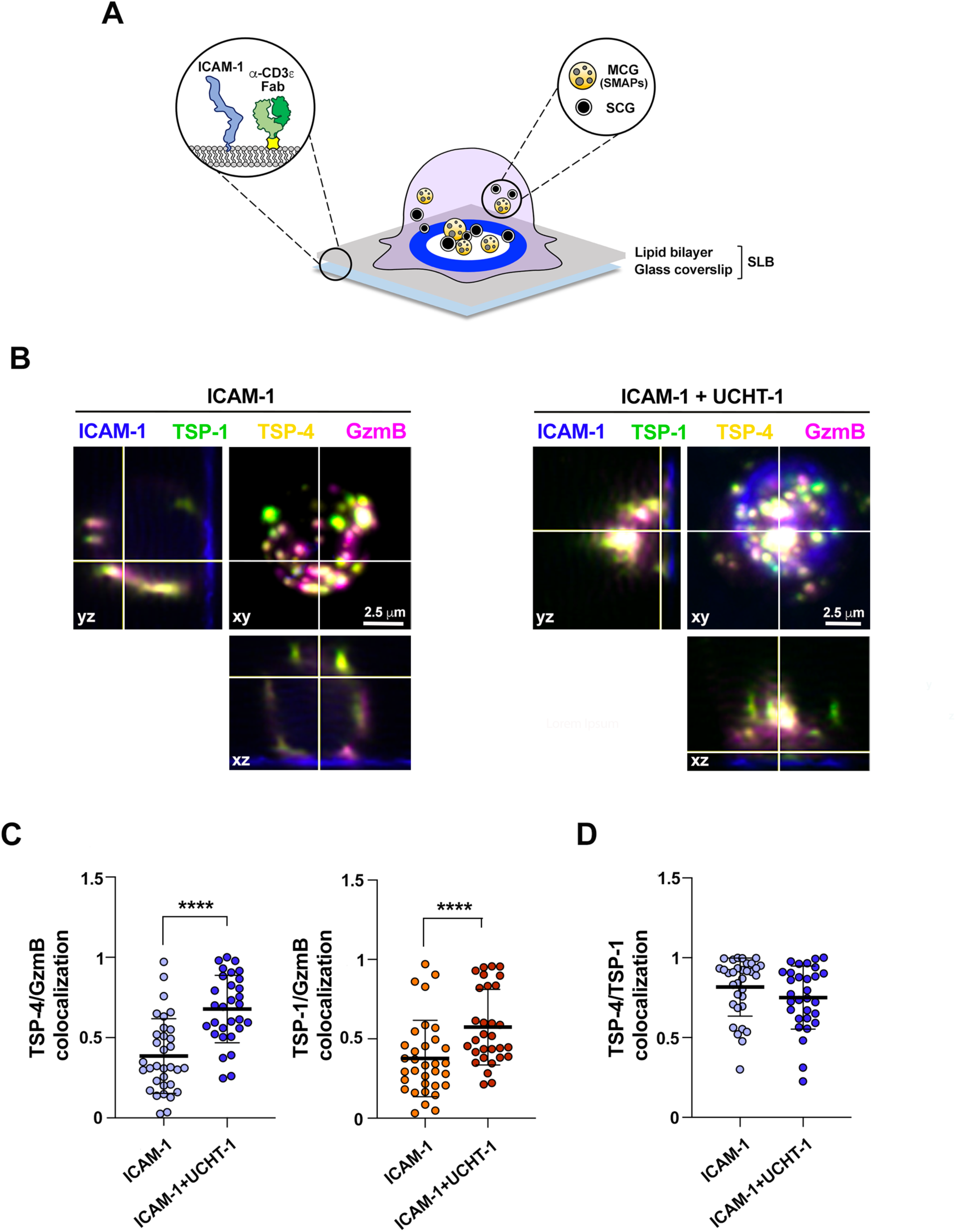
TSP-1 and TSP-4-containing LGs concentrate at the IS. (**A**) Schematic representation of the IS formed between a CTL and PSLBs presenting laterally mobile ICAM-1 and anti-CD3ε UCHT1 Fab’. The architecture of a mature IS is characterized by an ICAM-1-enriched ring (blue) surrounding an inner secretory domain, where multicore granules (MCG) containing SMAPs and single core granules (SCG) are focally released. (**B**) Representative maximum intensity z-projection and orthogonal views from confocal z-stacks of CTLs co-expressing either TSP-4-mCherry (yellow) and TSP-1-GFPSpark (green), and interacting with either non-activating [ICAM1-AF405] (*left*) or activating [ICAM1-AF405 + anti-CD3ε UCHT1 (unlabelled)] ligands (*right*) for 30 min. After fixation, cells were permeabilized and co-stained with anti-GzmB (magenta) antibodies. The formation of a mature IS is indicated by the presence of an ICAM-1 ring (blue). Scale bar: 5 μm. (**C,D**) Quantifications (mean±SEMSD) of the weighted colocalization using the Manders’ overlap coefficient (MOC) between each TSP signal and GzmB staining (C), and between TSP-4 and TSP-1 signals (D). Ndonor = 3, n ≥ 10 cells, Mann-Whitney test; **** p ≤ 0.0001, only significant differences are shown. Each dot represents one cell. 3D reconstructions of representative z-stacks are shown in Supplementary Videos 1-2.

### TSP-1 and TSP-4 are co-released in association with SMAPs

The localization of TSP-4 in LGs/MCGs that accumulate at the IS suggests that it might be released in association with SMAPs. To enhance GzmB imaging following SMAP release we used CTLs transfected with the GzmB-mCherry-encoding construct. The correct localization of GzmB-mCherry was confirmed by co-staining with anti-GzmB antibodies (Fig.S4A). Additionally, the colocalization of GzmB-mCherry with TSP-4 and TSP-1 at the IS formed on activating PSLBs recapitulated the one observed using anti-GzmB antibodies (Fig.S4B,C). For these experiments a GFPSpark-tagged TSP-4 construct was used (Fig.S2A,C). Next, CTLs were co-transfected with the TSP-4-GFPSpark and GzmB-mCherry constructs and plated on activating PSLBs to induce SMAP release, after which cells were flushed out by gentle washing. This leaves the synaptic output, that includes the SMAPs, on the PSLBs (Balint et al. 2020). TIRF-based imaging showed a significant colocalization of TSP-4 and GzmB in particles of size compatible with SMAPs (Fig.7A). Similar results were obtained when CTLs were co-transfected with TSP-1-GFPSpark and GzmB-mCherry (Fig.7B). When CTLs were co-transfected the constructs encoding TSP-4-mCherry and TSP-1-GFPSpark, TSP-4 and TSP-1 were found to co-localize in individual particles, of size compatible with SMAPs, released on the activating PSLBs (Fig.7C).

**Figure 7.**
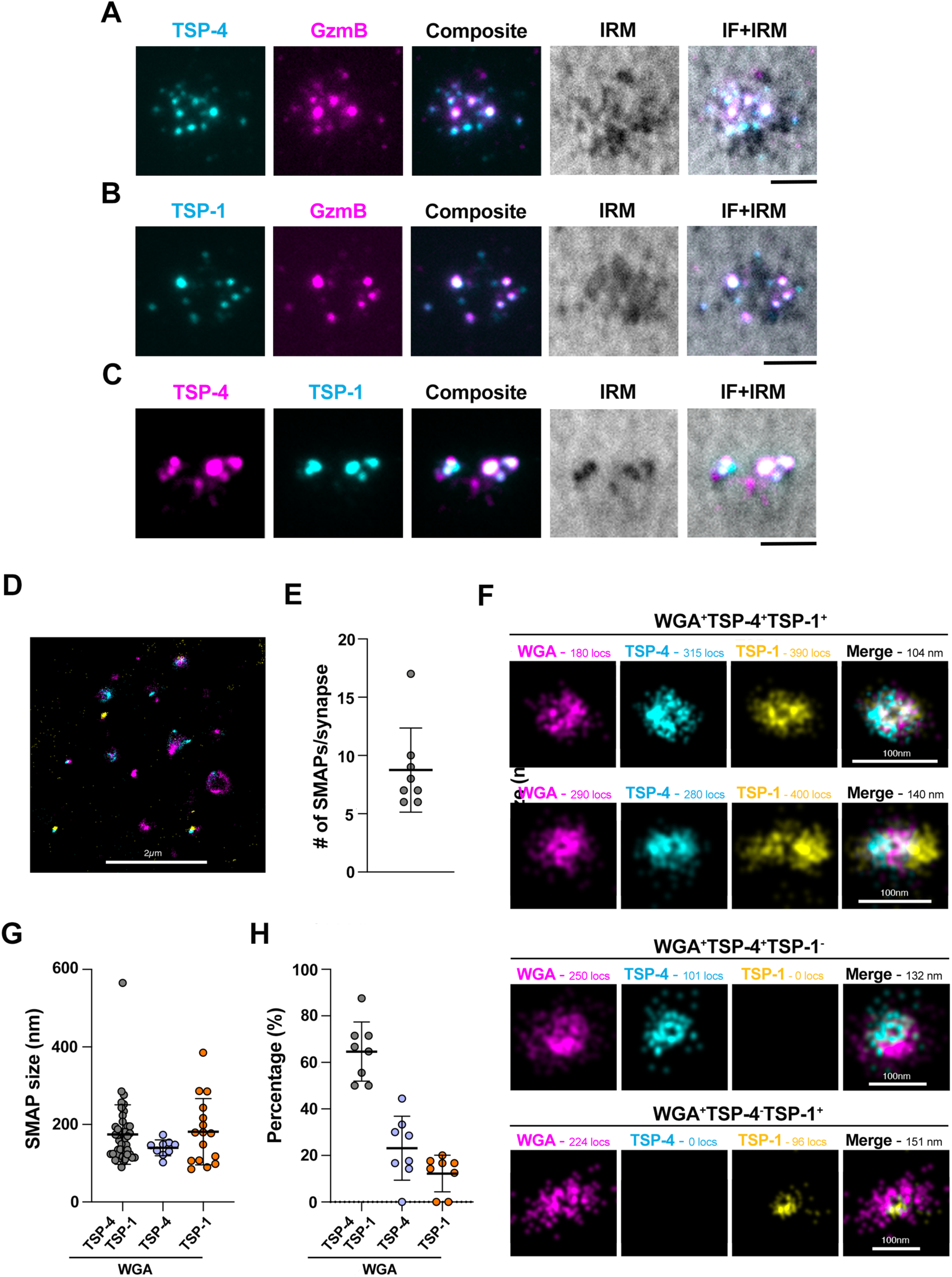
TSP-1 and TSP-4 are co-released at the IS in association with SMAPs. (**A**,**B**) TIRF images of TSP-4-GFPSpark^+^ (cyan) and TSP-1-GFPSpark^+^ (cyan) particles released by CTLs co-expressing fluorescently tagged TSP-4 (**A**) or TSP-1 (**B**) and GzmB-mCherry on activating PSLBs [ICAM-1 + anti-CD3ε UCHT1 Fab^’^ (unlabelled)]. (**C**) TIRF images of TSP-4-mCherry^+^ (magenta) and TSP-1-GFPSpark^+^ (cyan) particles released by CTLs expressing fluorescently tagged TSP-1 or TSP-4 on activating PSLBs [ICAM-1 + anti-CD3ε UCHT1 Fab’ (unlabelled)]. N_donors_ = 3 donors, n ≥ 10 cells. IRM, interference reflection microscopy. Scale bar: 2.5 μm. (**D**) Analysis of colocalization of TSP-1 and TSP-4 in SMAPs released by CTLs using 2D SMLM dSTORM to achieve single particle, single molecule resolution. The micrograph shows SMAPs deposited following IS formation of CTLs on the PSLB. The complete c-SMAC region is shown. Scale bar: 2 µm. (**E**) Average number of SMAPs per IS depicted as a box plot including outliers. **F**) Representative micrographs of single particles in CTL derived ISs. SMAPs are stained with WGA-AF647, TSP4-mCherry is probed with anti-RFP-AF555 and TSP1-GFPSpark is probed with anti-GFP-AF488. (**G**) Average size of SMAPs in each category depicted as a box plot including outliers. (**H**) Percentage of SMAPs positive for WGA, TSP1 and TSP4 derived from CTLs of 2 healthy donors (8 ISs and n=70 particles). Scale bars: 100 nm.

To better define the identity of these particles as SMAPs and visualize the respective localization of TSP-4 and TSP-1 therein we carried out a super-resolution analysis of the synaptic output released by CTLs on activating PSLBs by direct Stochastic Optical Reconstruction Microscopy (dSTORM). CTLs were co-transfected with the TSP-4-mCherry and TSP-1-GFPSpark constructs (transfection efficiency comparable, as assessed by flow cytometry; data not shown). The particles left on the PSLBs following CTL removal (8.8 particles/IS, Fig.7D,E) were stained with anti-GFP-Alexa488 and anti-RFP-Alexa555 for optimal blinking and WGA-Alexa647 to identify the SMAPs, and imaged by dSTORM (Fig.7D,F; Fig.S5; Fig.S6). As previously described for TSP-1 and WGA (Balint et al, 2020), TSP-4 displays ring staining consistent with its contributing to the glycoprotein shell of SMAPs (Fig 7F). The size of SMAPs released at the IS (Fig.7G) matched the size of MCG-associated SMAPs analyzed by CLEM (Fig.3E). A diffraction limited single particle colocalization analysis, performed to increase the number of observations, showed a prevalence of TSP1^+^TSP4^+^WGA^+^ particles as compared to TSP1^+^WGA^+^ or TSP4^+^WGA^+^ ones, consistent with the CLEM and STED analysis of SMAPs within MCGs (Fig.7H). Among the SMAPs single positive for either TSP released by cells co-expressing TSP-1-GFPSpark and TSP-4-mCherry, only a minor proportion were TSP1^+^WGA^+^ (Fig.7H). Together, the data show TSP-4 is part of the SMAP shell and that TSP-4 and TSP-1 are co-assembled and co-released in individual SMAPs.

### TSP-4 is required for CTL- and SMAP-mediated killing

While the specific role of SMAPs in CTL-mediated cytotoxicity is as yet not fully understood, TSP-1 deficiency in CTLs has been associated to impaired killing, although the contribution of SMAP-associated TSP-1 to this process has not been addressed (Balint et al, 2020). To assess the impact of TSP-4 deficiency on the killing ability of CTLs, CTLs transfected with TSP-4-targeting siRNAs or control scrambled siRNAs (Fig.8A) were mixed with SAg-pulsed target cells and cytotoxicity was tested by fluorimetry using the calcein release-based assay (Kummerow et al., 2014). TSP-4 knockdown (KD) CTLs showed an impaired ability to kill target cells, similar to TSP-1 KD CTLs (Fig.8B,C), indicating that TSP-4 is required for CTL-mediated killing.

**Figure 8.**
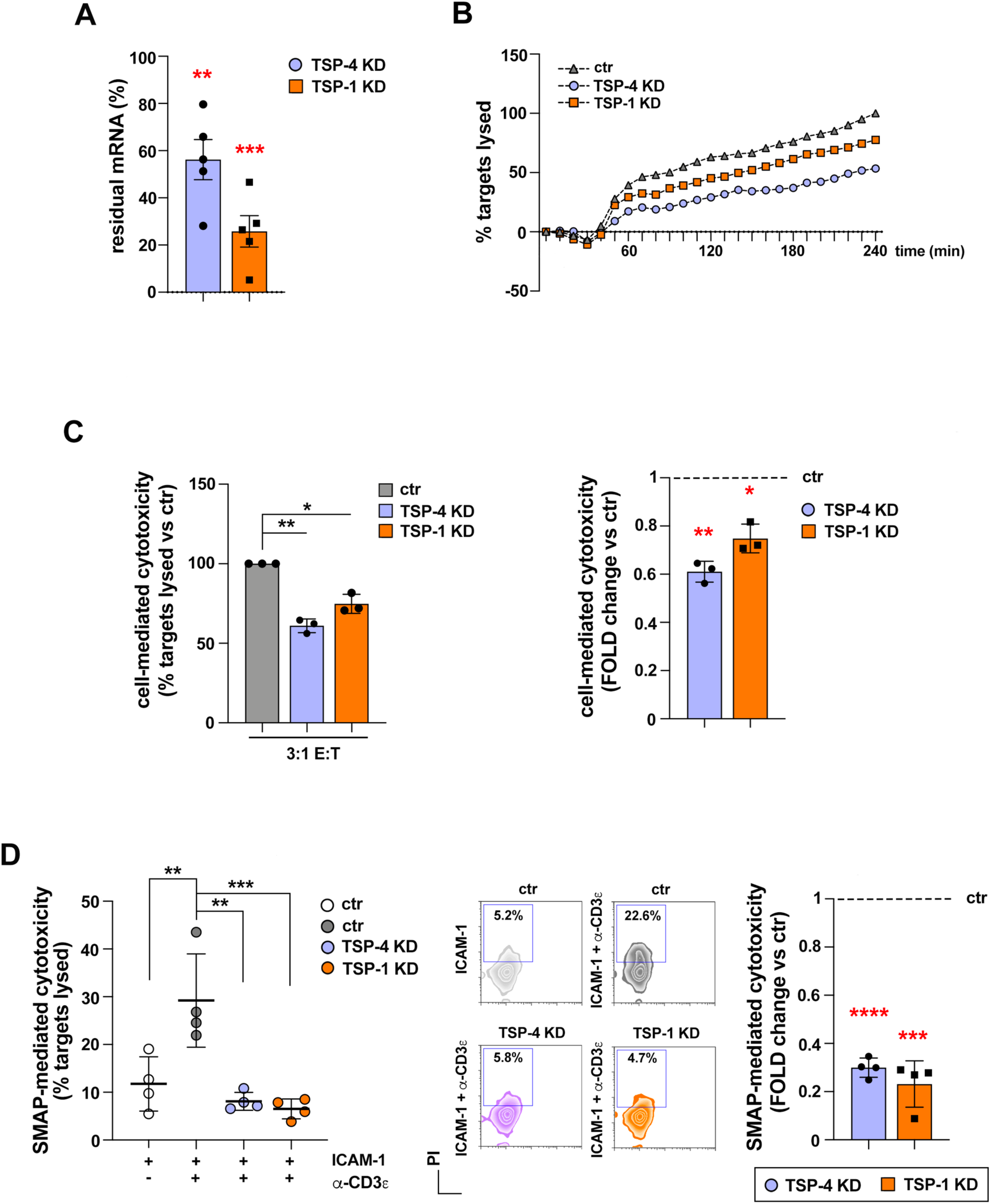
TSP-4 is required for CTL- and SMAP-mediated cytotoxicity. **(A)** RT-qPCR of TSP-4 and TSP-1 mRNA in control, TSP-4 KD and TSP-1 KD CTLs. Data are expressed (mean±SD) as residual % of mRNA in KD samples compared to control. N_donors_ = 3, one-sample t test; ** p ≤ 0.01, * p ≤ 0.05. (**B,C**) Fluorimetric analysis of cytotoxicity mediated by control (ctr), TSP-4 KD CTLs using the calcein release-based assay. Representative curves showing the kinetics of target cell death by CTLs at the effector:target ratio of 3:1 (**B**). Normalized quantification (mean±SD, ctr value = 100) of the target cell lysis (%) after 4 h (**C**, left) and cell-mediated cytotoxicity expressed as fold change in KD samples versus ctr. N_donors_ = 3, one sample t-test; ** p ≤ 0.01, * p ≤ 0.05, only significant differences are shown. (**D**) Flow cytometric analysis (mean±SD) of cytotoxicity mediated by the synaptic output of control (ctr), TSP-4 KD and TSP-1 KD CTLs plated on immobilized ICAM-1 or ICAM-1 + anti-CD3ε mAb. Quantification (mean±SD) of target cell lysis (%) (left), representative flow cytometry dot plots (middle) and SMAP-mediated cytotoxicity expressed as fold change in KD samples versus ctr (right). N_donors_ = 4, one-way ANOVA test (left) and one sample t test (right); **** p ≤ 0.0001, *** p ≤ 0.001, ** p ≤ 0.01, only significant differences are shown.

To elucidate whether this function of TSP-4 is mediated by SMAPs, SMAPs released by control and TSP-4 deficient CTLs were recovered on glass surfaces coated with ICAM-1 and activating anti-CD3 antibodies. Glass surfaces coated with non-activating ICAM-1 were used as negative control. MEC1 cells were then plated on the SMAPs captured on activating surfaces and incubated for 16 h, then recovered, stained with propidium iodide and analyzed by flow cytometry. The results show that SMAP-mediated cytotoxicity was impaired when SMAPs were released by TSP-4 KD CTLs (Fig.8D). This result was confirmed for SMAPs released by CTLs depleted of TSP-4 by CRISPR/Cas9 gene editing (Fig.S7). The analysis was extended to SMAPs released by TSP-1 KD CTLs. A shown in figure 8D, TSP-1 deficiency impaired SMAP-mediated killing. The data provide evidence that both TSP-4 and TSP-1 are required for the killing ability of SMAPs.

### Impaired cytotoxicity of CLL-conditioned CTLs is associated with impaired SMAP-mediated killing

We have previously reported that leukemic cells from CLL patients shape the lymphoid microenvironment to promote their survival while suppressing anti-cancer immunity through both contact-dependent and contact-independent interactions (Patrussi et al., 2021; Boncompagni et al., 2024). In particular, we showed that CLL cells enhance their surface expression of the immunosuppressive ligand PD-L-1 (Lopresti et al., 2024) and induce the upregulation of the inhibitory receptor PD-1 on CTLs (Boncompagni et al, 2024), leading to impaired CTL IS formation and CTL-mediated killing. We hypothesized that CLL cells could also suppress CTL function directly by tuning down the expression of the cytotoxic LG effectors. To test this hypothesis CD8^+^ T cells purified from healthy donors were differentiated to CTLs in the presence of culture supernatants from leukemic cells from CLL patients, using culture supernatants from healthy primary B cells as control (Fig.9A). As reported (Boncompagni et al, 2024), CTLs generated in leukemic cell-conditioned media show an impaired ability to kill CFSE-stained target cells following activation in the presence of SAgs as assessed by flow cytometry, gating on CFSE-positive (CFSE^+^) cells and using propidium iodide to identify dead cells (Fig.9B,C; Fig. S8 for gating strategy). RT-qPCR analysis of relative expression of shared SCG/MCG components (GzmA, GzmB, Prf, granulysin, serglycin) and of specific MCG/SMAP components (TSP-1, TSP-4) showed a reduction in the expression of GzmA, GzmB, serglycin and TSP-4 (Fig.9D). Notably, TSP-1 mRNA levels were not affected under these conditions (Fig.9D). The results indicate that CLL cells disable CTLs not only through the PD-1-dependent impairment of IS assembly and function, but also by interfering with LG biogenesis.

**Figure 9.**
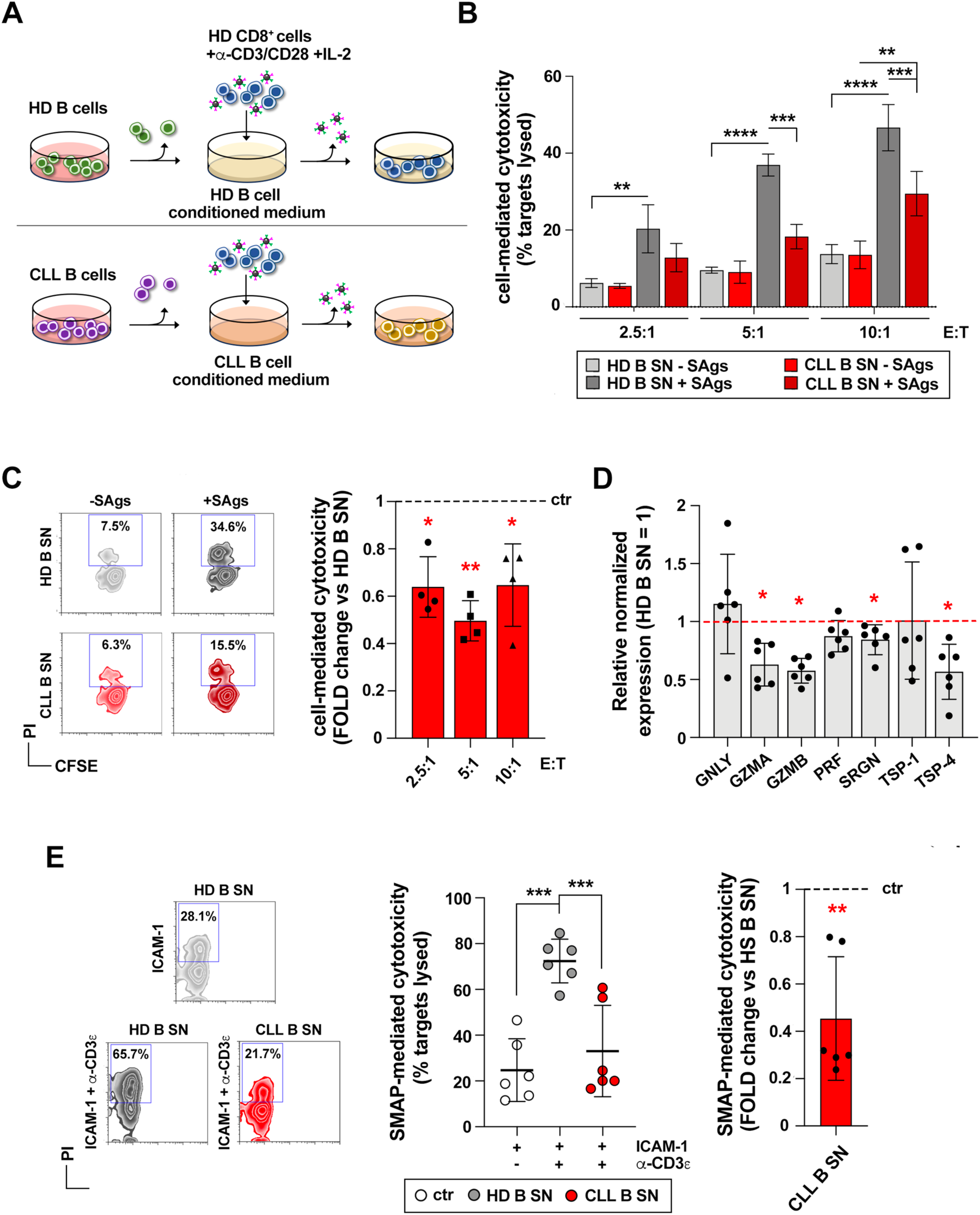
Impaired cytotoxicity of CLL-conditioned CTLs is associated with impaired expression of cytotoxic effectors and TSP-4 as well as SMAP-mediated killing. (**A**) Workflow for *in vitro* generation of CTLs from primary CD8^+^ T lymphocytes purified from buffy coats of healthy donors in the presence of conditioned media. (**B,C**) Flow cytometric analysis of cell-mediated cytotoxicity by CTLs generated from primary CD8^+^ T lymphocytes in the presence of media conditioned by either healthy B cells (HD B SN) or B cells purified from CLL patients (CLL B SN), and incubated with SAgs-pulsed CFSE-stained MEC1 B cells, used as targets, at the indicated E:T ratios. Cytotoxicity was assessed by using propidium iodide (PI) uptake by target cells. Quantification (mean±SD) of CFSE^+^/PI^+^ cells (%) (**B**), representative flow cytometry dot plots (**C**, left) and cell-mediated cytotoxicity expressed as fold change (**C**, right) in samples treated with CLL B SN vs samples treated with HD B SN. N_donors_ = 3, two-way ANOVA test (left) and one-sample t test (right); **** p ≤ 0.0001, *** p ≤ 0.001, ** p ≤ 0.01, * p ≤ 0.05, only significant differences are shown. (**D**) RT-qPCR of GZMA, GZMB, PRF, TSP-1, TSP-4, SRGN and GNLY mRNA in CTLs generated in the presence of media conditioned by either healthy B cells (HD B SN) or B cells purified from CLL patients (CLL B SN). The graph shows the normalized relative abundance of the transcripts (mean±SD, ctr value = 1). N_donors_ = 6, unpaired Student’s t-test; ** p ≤ 0.01, * p≤ 0.05, only significant differences are shown. (**E**) Flow cytometric analysis (mean±SD) of cytotoxicity mediated by the synaptic output of CTLs, generated in the presence of media conditioned by either healthy B cells (HD B SN) or B cells purified from CLL patients (CLL B SN) and plated on immobilized ICAM-1 or ICAM-1 + anti-CD3ε mAb surfaces. Quantification (mean±SD) of target cell lysis (%) (left), representative flow cytometry dot plots (middle; effector:target ratio 10:1 is shown) and SMAP-mediated cytotoxicity expressed as fold change (right) in samples treated with CLL B SN vs samples treated with HD B SN. N_donors_ = 3 CD8^+^ samples from healthy donors treated with either 6 HD B SN or 6 CLL B SN, one-way ANOVA test (left) and one-sample t test (right); *** p ≤ 0.001, ** p ≤ 0.01, only significant differences are shown.

The reduction of TSP-4 expression as well as of lytic effectors in CLL cell-conditioned CTLs suggests that the killing ability of SMAPs might be impaired. To test this, conditioned CTLs were plated on activating surfaces to capture the SMAPs. Killing by the SMAPs released by CLL cell-conditioned CTLs was reduced as compared to SMAPs derived from healthy B cell-conditioned CTLs, as assessed by plating MEC1 cells on the captured SMAPs followed by flow cytometric analysis of recovered cells stained with propidium iodide (Fig.9E). The results indicate that the SMAP biogenesis program of CTL is targeted by CLL cells for protection from CTL-mediated killing and may support the relevance of TSP-4 to the cytotoxic function of SMAPs.

## Discussion

SMAPs have been identified as a crucial element contributing to CTL and NK cell cytotoxicity (Balint et al., 2020; Ambrose et al, 2020). However, how SMAPs are assembled, how they are released and how they are taken up by target cells remains to be elucidated. The requirement for TSP-1 for CTL-mediated killing (Balint et al., 2020) suggests that components of the glycoprotein shell participate in SMAP function. Among the SMAP glycoproteins identified by mass spectrometry, only TSP-1 and Galectin-1 have been investigated to date, however the latter appears dispensable for the cytotoxic activity of CTLs (Balint et al., 2020). Here we have investigated the expression and function of TSP-4, which is among the glycoproteins picked up by the mass spectrometry analysis (Balint et al., 2020), and its structural similarity with TSP-1. We show that expression of TSP-4 and TSP-1 is regulated in a reciprocal manner during CD8^+^ T cell differentiation to CTLs. TSP-4 co-localizes with TSP-1 in a population of LGs that corresponds to the MCGs and that accumulates at the IS, where TSP-4 is co-released with TSP-1 in association with biologically active SMAPs. We show that TSP-4 as well as TSP-1 are required for the killing activity of SMAPs. Additionally, we provide evidence that CLL cells target SMAP biogenesis as a means to suppress CTL activity.

Expression of TSPs has been documented in a variety of cell types and tissues both in physiological and pathological conditions. While largely ubiquitous, expression of individual TSPs is spatiotemporally regulated, likely reflecting the requirement for these proteins in specific cellular processes (Raugi et al., 1987; Stenina et al., 2003; Dabir et al., 2008; Stenina-Adognravi, 2014). Here we provide the first evidence that TSP-4 is expressed in CD8^+^ T cells throughout their differentiation to cytotoxic effectors. Interestingly, TSP-1 and TSP-4 show an opposite expression pattern during CD8^+^ T cell differentiation to CTLs, with a progressive increase in TSP-4 paralleled by a concomitant drop in TSP-1. Beyond their involvement in SMAP biogenesis and function, it remains to be elucidated whether TSP-1 and TSP-4 play distinct roles at other stages of CTL differentiation or in other effector functions. Alternatively, TSP-4 could take over the function of TSP-1 as expression of the latter declines. It is noteworthy that a similar opposite co-regulation of TSP-1 and TSP-4 expression has emerged from the analysis of gene expression databases in brain and breast cancer (Sorlie et al., 2001; Bredel et al., 2005; Liang et al., 2005; Sun et al., 2006; Turashvili et al., 2007; Ma et al., 2009). Our findings show that this can also occur in physiological conditions, with potential relevance to other cell types.

Our data show that under homeostatic conditions TSP-1 and TSP-4 are pre-assembled in GzmB^+^LAMP-1^+^ vesicles. While the vesicular trafficking pathway that regulates their transport to these LGs remains to be characterized, only a proportion of GzmB^+^ vesicles are also TSP-1^+^ or TSP4^+^, consistent with the existence of two classes of LGs, the SCGs and MCGs, that differ for the presence of TSP-1 (Chang et al, 2022) and TSP-4 (this report), among other differences. This indicates a bifurcation of the GzmB trafficking pathway, with one branch leading to SCGs and another intersecting with TSP-1/4 trafficking at MCGs. Interestingly, our observation that TSP4^+^TSP1^+^ and TSP4^+^ MCGs are more prevalent than TSP1^+^ MCGs, highlighted by the CLEM and STED analysis and reflected at the SMAP level by the dSTORM analysis, suggested the hypothesis that TSP-1 may be incorporated into SMAPs mainly after TSP-4. Our finding, based on the RUSH experiments, that TSP-4 localizes indeed to LGs prior to TSP-1 and that, additionally, it is required for LG localization of TSP-1, indicates a diversification within the TSP-1/4 trafficking pathway and supports a tight, non-redundant interplay of the two TSPs in the stepwise process of SMAP biogenesis. It is noteworthy that the colocalization of both TSP-4 and TSP-1 with GzmB increases after activation, as observed in the PSLB experiments, suggesting a maturation step involving vesicle fusion triggered by CTL recognition of its cognate target leading to the release of fully armed SMAPs. Vesicle fusion events during LG maturation have been reported previously for endosomes and SCGs (Ménanger et al., 2007; Sanchez-Ruiz et al., 2011), suggesting that similar mechanisms might apply to MCGs.

Using activating PSLBs, we found that the pre-formed TSP-1^+^/TSP-4^+^ LGs accumulate at the cSMAC during IS assembly and that TSP-1 and TSP-4 are co-released as SMAPs endowed of killing activity. Owing to the lack of antibodies suitable for imaging, these observations were carried out using fluorescently tagged TSP-1 and TSP-4. While this provides information on the relative localization of the two TSPs within CTLs prior and following stimulation, they do not answer the issue of the relative SMAP content of endogenous TSP-4 vs TSP-1. The mass spectrometry analysis captured a substantially larger abundance of TSP-1 vs TSP-4 peptides (Balint et al., 2020), suggesting that, despite the downregulation in TSP-1 expression, the residual amounts are sufficiently high to play a role in SMAPs and, accordingly, TSP-1 KO was shown to impair the killing ability of CTLs (Balint et al., 2020) and of SMAPs (this report). Here we show that TSP-4 deficiency leads to defects both in CTL- and SMAP-mediated killing, indicating that, despite the fact that we do not have sufficient information as to the relative amounts of endogenous TSP-4 vs TSP-1, both TSPs participate in SMAP biogenesis and function. *Thbs1*^-/-^ and *Thbs4*^-/-^ mice may help unravel this issue. Both mice are viable but display a number of developmental and functional defects, from major lung abnormalities associated with pneumonia in *Thsp1*^-^/- mice (Lawler et al., 1998) to neuronal and cardiac defects in *Thsp*4^-/-^ mice (Girard et al., 2014; Frolova et al., 2012; Palao et al., 2018). Immune functions of these mice have not been examined to date, although an increase in white blood cell numbers, including lymphocytes, has been observed in *Thsp1*^-/-^ mice (Lawler et al., 1998). Elucidating the role of TSP-1 and TSP-4 function in CTLs and NK cells in the in vivo setting of *Thsp1*^-/-^ and *Thsp4*^-/-^ mice may provide key information not only on their respective role in the assembly of SMAPs but also on the specific role of the SMAP pathway in CTL-mediated cytotoxicity towards specific pathogens or roles in immune homeostasis. It should be pointed out that, while both TSP-1 and TSP-4 are required for the killing activity of SMAPs, they might participate in CTL-mediated killing by affecting also other key features of the killing process, one of these being CTL adhesion to target cells, that TSPs might affect through binding to integrins and other adhesion molecules (Adams and Lawler, 2011).

Tumor cells have evolved a variety of strategies to escape elimination by CTLs, including preventing their recruitment, enhancing the generation and function of Tregs by promoting the expression of inhibitory checkpoints and inhibiting the assembly of functional immune synapses, as exemplified by CLL (Gribben et al., 2008; Nicholas et al., 2016; Arruga et al., 2020; Apollonio et al., 2021). We have previously reported that CLL cells have the ability to rewire cellular components of the lymphoid niche, which is their primary TME, to enhance their homing to and persistence therein, where they are exposed to survival factors and protected from apoptosis-inducing drugs. These include stromal cells, which upregulate their production of homing chemokines (Patrussi et al., 2021), and CTLs themselves, which upregulate PD-1 expression (Boncompagni et al., 2024). Here we identified a new, additional immunosuppressive strategy whereby CLL cells disable CTLs by downregulating the expression of key components of their cytotoxic machinery. Interestingly, these include not only cytotoxic molecules shared by SCGs and MCGs, but also TSP-4. Taken together with the killing defect of SMAPs released by TSP-4 deficient CTLs, the fact that SMAPs released by CTLs exposed to the culture supernatants of CLL cells have an impaired killing ability further highlights the importance of TSP-4 in SMAP function, although further work is required to determine if SMAPs play an important role in killing of CLL cells.

In conclusion, our data identify TSP-4 as a new participant in SMAP biogenesis and function. This not only advances our knowledge of the mechanisms underpinning CTL-mediated killing but also potentially paves the way to new, SMAP-based, cell-free therapeutic strategies through engineering TSPs to remove the molecular determinants responsible for non-selective cell adhesion, while endowing them with tumor antigen specificity.

## Materials and Methods

### Ethics

Either buffy coats (Siena University Hospital) or leukocyte reduction system chambers, a by-product of platelet collection from healthy blood donors (Institute of Clinical Hemostaseology and Transfusion Medicine, Saarland University Medical Center; Oxford) were obtained from the local blood banks. All healthy donors and CLL patients provided voluntary informed written consent to use their blood for research purposes. All procedures were performed according to the Declaration of Helsinki guidelines. Research with human PBMC and CLL-derived samples has been approved by the local ethic committee of Siena University (20759; Prof. Baldari), Saarland University (98/15; Dr. Krause), Oxford University (REC 11/H0711/7; Prof. Dustin) and Padua University (23229; Prof. Trentin).

### CLL patients

Peripheral blood samples were collected from 16 previously untreated CLL patients. Diagnosis of CLL was made according to international workshop on CLL (iwCLL) 2008 criteria (Hallek et al Blood 2018). The immunophenotypic analysis of lymphocytes obtained from peripheral blood of CLL patients was performed as reported (Patrussi 2021 Blood). B cells from 16 buffy coats, used as healthy population controls, were purified by negative selection using RosetteSep B-cell enrichment Cocktail (StemCell Technologies, Vancouver, Canada) followed by density gradient centrifugation on Lympholite (Cedarlane Laboratories, The Netherlands), as reported (Patrussi 2021 Blood).

### Plasmids and antibodies

Expression constructs for human TSP-4 tagged with fluorescent tags were generated by insertion of their respective coding sequences into a pMax backbone. GFPSpark and mCherry cDNAs were amplified using the primers listed in Table S2, and pCMV-GzmB-mC (Ming et al., 2015) and pCMV3-TSP1-GFPSpark (Bálint et al., 2020) as templates, then inserted into the pMax via restriction digestion (Thermo Fisher Scientific) and ligation by T4 DNA ligase (Promega Corporation). TSP-4 cDNA was amplified from pRP[Exp]-mCherry/Neo-CAG>hTHBS4[NM_003248.6] (ID VB900137-1200xfw, VectorBuilder) removing the terminal STOP codon by using primers listed in Table S2 and inserted into the pMax-GFPSpark and pMax-mCherry vectors using KpnI and EcoRI restriction sites. The pMax-GFPSpark and pMax-mCherry vectors (empty vectors) were used as negative controls. The TSP-1-GFPSpark coding sequence was subcloned from pCMV3-TSP1-GFPSpark (Bálint et al., 2020) to a pMax vector using KpnI and XbaI restriction sites. The constructs were confirmed by sequencing. The human GzmB-mCherry construct was kindly provided by Prof. Jens Rettig (Ming et al., 2015).

To generate pMax-TSP1-linker-3xFLAG, the TSP1 gene sequence was amplified from the background vector pMax-TSP1-GFPSpark using the primers listed in Table S2 to add KpnI and XbaI sites at the 5’ and 3’ ends. The linker and the 3xFLAG were created through the annealed oligo cloning method, using the primers listed in Table S2. The oligos were mixed in equal molar amounts, heated to 95°C for 5 minutes, and gradually cooled to room temperature. The annealed linker was digested with XbaI and NheI, and 3xFLAG with NheI and XhoI. Subsequently, the background vector plasmid pMax was digested with KpnI and XhoI restriction enzymes. The digested products were then ligated into the pMax-digested vector, creating the pMax-TSP1-linker-3xFLAG vector.

Cloning of pMax-TSP4-linker-3xHA plasmid was achieved by first digesting the pMax-THBS4-mCherry vector with EcoRI and XhoI restriction sites to remove the mCherry tag. Then, the linker was generated through the annealed oligo cloning method, using the primers listed in Table S2. The annealed linker was digested with EcoRI and XbaI. The HA tag was digested from the pMax-3xFLAG-mFwe 2-3xHA vector using XbaI and XhoI restriction enzymes. The digested linker and HA tag were then ligated with the pMAX-THBS4-digested vector to obtain the final clone. The final vectors were confirmed by sequencing the plasmids using the respective forward and reverse primers, a process conducted by Microsynth Seqlab.

To clone the reporter of interest (i.e. TSP-4 and TSP-1) into the RUSH plasmids, the sequence encoding human TSP-1 fused to a streptavidin binding peptide (SBP) was synthetized in vitro (ID VB210412-1326qnm, VectorBuilder) and AscI and SbfI sites were added at the 5’ and 3’ ends. Both insert and the Ii-Str_TNF-SBP-EGFP (#65280, Addgene) vector were digested using AscI and SbfI (New England Biolabs) and ligated by T4 DNA ligase (Promega Corporation). Human TSP-4 cDNA was amplified from pRP[Exp]-mCherry/Neo-CAG>hTHBS4[NM_003248.6] (ID VB900137-1200xfw, VectorBuilder) by using the primers listed in Table S2 and inserted into the Str-KDEL_TNF-SBP-mCherry (#65279, Addgene) via restriction digestion with AscI and EcoRI (Thermo Fisher Scientific) and ligation by T4 DNA ligase (Promega Corporation). The constructs were confirmed by sequencing.

Primary commercial antibodies used in this work and their application are listed in Table S1. The bacterial superantigens (SAgs) staphylococcal enterotoxins A (SEA), B (SEB) and E (SEE) were purchased from Toxin Technology Inc. Bovine serum albumin (BSA) heat shock fraction (pH 7), poly-L-lysine, propidium iodide and saponin were obtained from Merck, Triton X-100 and BSA fraction V (pH 7) from PanReac Applichem.

### RNA purification and RT-qPCR

RNA was extracted from 2-5×10^6^ freshly isolated CD8^+^ T cells (day 0) and CTLs (day 5 and 7) and from 0.3-0.5×10^6^ CTLs transfected with TSP-1/4-targeting siRNAs or control scrambled siRNA using the RNeasy Plus Mini Kit, reverse transcribed to first-strand cDNA iScript cDNA Synthesis Kit, and analyzed by quantitative PCR (qPCR) by using the SSo Fast™ Eva Green® Super Mix (Bio-Rad) and the CFX96 Real-Time PCR Detection System (Bio-Rad). After an initial denaturation at 95°C for 3 min, samples were subjected to 42 cycles of denaturation at 95°C for 10 sec and primer annealing plus extension at 60°C for 30 sec, followed by 10 sec at 95°C and a denaturation gradient from 65°C to 95°C with 0.5°C increment every 5 sec to generate melt curves. Results were processed and analyzed using the CFX Manager Version 1.5 software (Bio-Rad). The abundance of GLNY, GZMA, GZMB, PRF, SRGN, TSP-1 and TSP-4 transcripts was determined on duplicate samples using the ΔΔCt method (Livak et al., 2001, Pfaffl et al., 2001) and normalized to 18S ribosomal RNA (Fig. 1A,8C) or to HPRT-1 RNA (Fig. 7A). Specific primers used to amplify cDNA fragments corresponding to human transcripts are listed in Table S2.

### Cell lysis and immunoblots

Primary CD8^+^ T cells were washed twice with ice-cold PBS 1X and lysed in 20 mM Tris-HCl pH8, 150 mM NaCl, 1% Triton X-100 in the presence of Protease Inhibitor Cocktail Set III (Calbiochem®) and 0.2 mg sodium orthovanadate for 5 min on ice. The homogenates were centrifuged at 16,100 x g for 20 min at 4°C and the soluble fractions were recovered. Protein concentration in post-nuclear supernatants was determined using the Quantum Protein Bicinchoninic Acid (BCA) Assay (Euroclone) and the iMark™ microplate reader (Bio-Rad) to measure the absorbance at 570 nm. 10 µg of protein extract (post-nuclear supernatant) were denatured in reduced 1X Bolt™ LDS Sample Buffer (Invitrogen™) for 5 min at 100°C and resolved on precast polyacrylamide gradient gels Bolt™ 4 to 12%, Bis-Tris, 1.0 mm, Mini Protein Gels (Invitrogen™) in 1X Bolt™ MES SDS Running Buffer with a constant voltage of 150 V. Separated proteins were transferred to nitrocellulose membranes (Amersham™ Protran®) in 20 mM Tris, 0.2 M glycine, 20% (v/v) ethanol for 75 min setting a constant current of 250 mA. Membranes were blocked in 4% (w/v) non-fat dry milk in PBS 1X (140 mM NaCl, 2.7 mM KCl, 10 mM phosphate buffer pH 7.4) + 0.02% (v/v) Tween-20 (PBS-T), probed with primary abs (Table S1), then incubated with horseradish peroxidase-labeled goat anti-mouse or anti-rabbit secondary IgG (heavy and light chains) (Jackson ImmunoResearch Inc) (Table S2) and revealed using the chemiluminescent detection kit SuperSignal™ West Pico PLUS Chemilumiscent Substrate (Thermo Fisher Scientific). Chemiluminescence signals were acquired by using Alliance Q9-ATOM imaging system (UVITEC, Cambridge, UK) and NineAlliance x64 software (UVITEC). The densitometric analysis of protein bands was carried out using ImageJ (National Institutes of Health, USA).

### Flow cytometry

Flow cytometry analysis of surface CD62L and CD45RA was carried by incubating 0.2-0.15×10^6^ freshly isolated CD8^+^ T cells (day 0, not shown) and 5-day and 7-day (not shown) CTLs in the dark for 30 min on ice with APC-labeled anti-hCD62L and FITC-labeled anti-hCD45RA (Table S1). Viable cells were gated based on size and granularity. Sample acquisition was performed with a Guava® easyCyte™ Flow Cytometer (Merck Millipore) using the appropriate lasers and emission filters, and data were analyzed using FlowJo 6.1.1 software (TreeStar Inc.). Staining was performed in two independent experiments with cells from different donors. To determine the cells single positive for each marker and both double negative and double positive, the density plot was split into four quadrants and the relative abundance (%) of CD45RA and CD2L within viable cells identified CD45RA^+^/CD62L^+^ naïve cells, CD45RA^-^/CD62L^+^ central memory cells, CD45RA^-^/CD62L^-^ effector memory cells and CD45RA^+^/CD62L^-^ terminal effector memory re-expressing CD45RA (TEMRA) cells in the samples was then quantified.

For flow cytometry analysis of CTLs transiently transfected with the constructs for TSP-1-GFPSpark, TSP-4GFPSpark and TSP-4-mCherry (alone or in combination), 24 h after transfection 0.2×10^6^ cells were fixed with Cyto-Fast Fix/Perm Solution (#750000133, BioLegend) in the dark for 15 min at room temperature and labelled with anti-GFP and anti-RFP primary antibodies in Cyto-Fast Perm/Wash solution 1X (#750000135, BioLegend) for 30 min on ice following washing with Cyto-Fast Perm/Wash solution 1X and labelling with AF488-anti-mouse or AF647-anti-rabbit antibodies in Cyto-Fast Perm/Wash solution 1C for 30 min on ice in the dark. Untransfected cells and cells with only secondary antibodies were used as negative controls. Transfection efficiency was calculated as the percentage of cells showing above-background levels of fluorescence. In double transfectants transfection efficiency was comparable for the two constructs in all experiments.

Flow cytometry was perfomed using a Guava® easyCyte™ Flow Cytometer and guavaSoft InCyte 2.7 software (Merck Millipore) using the appropriate lasers and emission filters. Data were analyzed using guavaSoft InCyte 2.7 (Merck Millipore) and FlowJo version 6.1.1 (Tree Star Inc.).

### Cell culture and transfection of CTLs for confocal imaging

Primary human CD8^+^ T lymphocytes were isolated from buffy coats of anonymous healthy donors and purified (>95% purity) by negative selection using the RosetteSep™ Human CD8+ T Cell Enrichment Cocktail (STEMCELL Technologies). Buffy coats were collected at the Siena University Hospital after obtaining written informed consent from each donor and processed following standard ethical procedures outlined in the Declaration of Helsinki. Freshly purified CD8^+^ T cells (day 0) were activated by adding Dynabeads™ Human T-Activator CD3/CD28 (Gibco™) at a cell/bead ratio of 1:0.5 for 48-64h and expanded in complete R10 medium [RPMI-1640 medium containing 20 mM HEPES and 2.05 mM L-glutamine (#7388, Merck) supplemented with 10% iron-enriched bovine calf serum (BCS; GE Healthcare HyClone), 120 µg/ml penicillin, 1% MEM Non-Essential Amino Acids (MEM NEAA; Gibco™) and 100 U/ml of recombinant human rhIL-2 (Miltenyi)] for further 3-5 days (days 5 and 7 after isolation) to generate CTLs as previously reported (Onnis et al., 2022).

CTLs were transiently transfected with the constructs pMax-TSP-1-GFPSpark, pMax-TSP-4-GFPSpark, pMax-TSP-4-mCherry, pMax-GzmB-mCherry constructs, and the GFPSpark and mCherry control vectors (DNA/cell ratio = 1.5 µg/10^6^ cells in single transfections and 1.2 µg/10^6^ cells in co-transfections) by using the Human T Cell Nucleofector Kit (Amaxa Biosystem) and the Amaxa Nucleofector II system (Lonza), Program T-023. Cells were cultured with 35 U/ml of rhIL-2 in complete R10 medium at a cell density of 1×10^6^ cells/ml and analyzed 24 h after transfection.

### Spinning Disk Confocal Microscopy imaging (SDCM)

3D confocal microscopy imaging was carried out on fixed samples at 200 nm steps using a Nikon ECLIPSE Ti2-E microscope equipped with a Yokogawa CSU-W1-SoRA spinning disk confocal unit (50 μm pinhole size) and a Photometrics Prime BSI (Nikon), and NIS-Elements AR 5.42.02 64-bit software. A 100X/1.4 NA oil immersion objective were used for image acquisition. For SDCM imaging, cells were plated on 10-well diagnostic microscope slides (Epredia) coated with 0.1% (w/v) poly-L-lysine (Merck) in H2O, fixed 4% PFA/PBS (v/v) for 15 min at room temperature. After fixation, cells were washed in PBS 1X (v/v), permeabilized and stained with primary abs (Table S2) in 1% (w/v) BSA, 0.1 (v/v) % Triton X-100 or saponin in PBS 1X over-night at 4°C. Samples were washed with PBS 1X (v/v) and incubated with fluorescent secondary abs (Table S2) for 45 min at room temperature. Samples were washed again with PBS 1X (v/v), mounted in 90% glycerol/PBS (v/v) and inspected by SDCM.

Z-stacks were denoised and deconvoluted using the Richardson-Lucy algorithm (Richardson et al., 1972; Lucy et al., 1974) and 10 iteractions. Post processing and analysis of fluorescence images was performed with Fiji (National Institute of Health). Colocalization analyses on median optical sections and 3D reconstructions was performed using JACop plug-in (Bolte and Cordelières, 2006) to calculate the Manders’ overlap coefficient (MOC). MOC varies from 0 to 1, with 0 standing for non-overlapping images and 1 for 100% colocalization between two images. Specifically, M1 is defined as the ratio of the “summed intensities of pixels from the green image for which the intensity in the red channel is above zero” to the “total intensity in the green channel” and M2 is defined conversely for red (Manders et al., 1992).

### PSLB preparation

PSLB were prepared as previously described (Demetriou et al., 2020; Balint et al., 2020). Small unilamellar vesicles composed of 0.4 mM solution of lipids in PBS with 75 mol% 1,2-dioleoyl-sn-glycero-3-phosphocholine supplemented (DOPC) and 25 mol% 1,2-dioleoyl-sn-glycero-3-[(N-(5-amino-1-carboxypentyl) iminodiacetic acid) succinyl] (DOGS-NTA) (Avanti Polar Lipids Inc.) were prepared by extrusion using the Avanti Miniextruder with a 100 nm filter. Protein concentrations required to achieve desired densities on bilayers were calculated from calibration curves that were constructed from flow cytometric measurements of fluorescent proteins attached on bilayers formed on glass beads, compared with reference molecules of equivalent soluble fluorophores (MESF)-beads (Bangs Laboratories).

Sticky 6-channel slides (sticky-Slide VI 0.4, Ibidi) were glued to cleanroom cleaned coverslips 0.170×0.005 mm thickness (SCHOTT MINIFAB Diagnostics – NEXTERION® Glass). A 1:1 mixture of DOGS-NTA lipids and DOPC stocks were loaded in each channel and incubated for 20 min to generate mobile PSLBs. Channels were washed twice with 0.1% (w/v) BSA in HEPES Buffered Saline 1X pH 7.2 (0.2 mM HEPES, 1.37 mM NaCl, 50 nM KCl, 7 nM Na2HPO4, 60 nM D-glucose, 1 mM CaCl2, 2 mM MgCl2) and blocked with 2% BSA/HBS (w/v) supplemented with 100 µM NiSO4 to generate Ni-chelating lipids that anchor His-tagged proteins. The channels were washed again twice with 0.1% BSA/HBS before adding 200 molecules/µm^2^ AlexaFluor™ 405-labeled ICAM-1 alone (steady-state conditions) or in combination with 30 molecules/µm^2^ unlabeled anti-CD3ε (UCHT1) Fab’ (activating conditions). Excess proteins were removed by washing twice with 0.1% BSA/HBS.

### IS formation on PSLBs

1×10^6^ CTLs were plated onto non-activating (ICAM-1) and activating (ICAM1 + anti-CD3ε Fab’) surfaces for 30 min at 37°C and 5% CO_2_, then fixed with 4% paraformaldehyde (PFA) in PBS 1X (v/v) for 15 min at room temperature. After fixation, samples were washed twice with 0.1% BSA/HBS, blocked and permeabilized with 1% (w/v) BSA, 0.1 (v/v) % Triton X-100 in PBS 1X for 1h at room temperature and then stained with primary abs (Table S2) over-night at 4°C. Samples were washed twice with 0.1% BSA/HBS and incubated with fluorescent secondary abs (Table S2) for 45 min at room temperature. Samples were washed again twice with 0.1% BSA/HBS and analyzed by SDCM.

### Total Internal Reflection Microscopy (TIRF) imaging of released SMAPs

1×10^6^ CTLs were plated onto non-activating (ICAM-1) and activating (ICAM1 + anti-CD3ε Fab’) PSLBs and incubated for 90 min at 37°C and 5% CO_2_, then flushed out by washing three times with ice-cold PBS. SMAPs released by CTLs and remained attached to PSLBs were fixed with 4% PBA/PBS (v/v) for 15 min at room temperature and analyzed by TIRF microscopy using a Nikon ECLIPSE Ti2-E microscope equipped with a Yokogawa CSU-W1-SoRA spinning disk confocal unit and a Photometrics Prime BSI (Nikon). A 100X/1.4 NA oil immersion objective was used for image acquisition.

### Cell culture and transfection of human CTLs for CLEM and STED microscopy

Human PBLs were obtained from healthy donors as previously described in Schwarz et al. (2007). Naïve CD8^+^ T cells were isolated using the Dynabeads^TM^ untouched human CD8 T-cell isolation kit (Invitrogen), stimulated with Dynabeads™ Human T-Activator CD3/CD28 (Gibco™) and cultured for 5 days in AIMV medium supplemented with 10% fetal calf serum (FCS) and 100 U/ml of rhIL-2 (Gibco). 8×10^6^ CTLs were transfected with 6 µg of plasmid DNA of each construct with P3 Primary Cell 4D-Nucleofector X Kit (Lonza). For the CLEM experiments the cells were transfected with TSP-1-GFPSpark and TSP-4-mCherry, while for the STED microscopy experiments the CTLs were transfected with TSP1-Flag and TSP4-HA (Alawar et al., 2024). Cells were washed 6 h after plasmid transfection and replated in fresh medium. 16 h after transfection, cells were either directly immune-stained for STED microscopy or incubated with WGA-AlexaFluor™ (AF) 647 (1:1000) for 2 hours prior the CLEM experiment. After two wash steps in ice cold RPMI-1640 medium, cells were collected in AIMV medium with 30% FCS and high pressure frozen.

### Post-embedding correlative fluorescence electron microscopy (CLEM)

Workflow is shown in Fig. S3. CTLs were seeded on sapphire discs coated with poly-L-ornithine (0.1 mg/ml) and anti-CD3 UCHT1 (30 µg/ml, BioLegend # 300402) in flat specimen carriers (Leica). After incubation at RT (20°C ± 2°C) for 15 min, samples were vitrified in a high-pressure freezing system (Leica EM PACT2). Samples were further processed in a freeze-substitution apparatus (AFS2; Leica) as described in Chang et al. (2022). Briefly, all samples were transferred to the precooled (−130°C) freeze-substitution chamber of the AFS2. The temperature was increased from −130 to −90°C over 2 h. Freeze substitution was performed from −90 to −70°C over 20 h in anhydrous acetone and from −70 to −60°C over 24 h with 0.3% (w/v) uranyl acetate in anhydrous acetone. At -60 °C samples were infiltrated with increasing concentrations (30, 60, and 100% for 1 h each) of Lowicryl (3:1 K11M/HM20 mixture; Electron Microscopy Sciences). After 5 h of 100% Lowicryl infiltration, samples were UV polymerized at −60°C for 24 h and for additional 15 h at a linear temperature increase to 5°C. Samples were stored in the dark at 4°C until further processing. After removal of the carriers and the sapphire discs ultrathin sections (100 nm) were cut using an UC7 (Leica) and collected on carbon-coated 200 mesh copper grids (Plano). 1 day after sectioning the grids were stained with DAPI for 3 min (1/1000), washed and sealed between two coverslips for high-resolution SIM imaging. The same grids previously analyzed with SIM were stained with uranyl acetate and lead citrate and recorded with the Tecnai 12 Biotwin electron microscope. Only CTLs with well-preserved membranes, cell organelles and nuclei were analyzed and used for correlation. For correlation, the DAPI 405 nm image showing the labeled nucleus of the cell was used to find the optimal overlay with the electron microscope images. The final alignment defines the position of the fluorescent signals within the cell of interest. The Images were overlaid in Corel DRAW 2021.

### Structured illumination microscopy (SIM)

For high-resolution SIM (ELYRA PS.1; Carl Zeiss Microscopy) of the ultrathin sections, images were acquired by using the 63X Plan-Apochromat (NA 1.4) objective with excitation light of 405-, 488-, 561- and 642-nm wavelengths to visualize DAPI, TSP-1-GFPspark, TSP-4-mCherry and WGA-AF647, respectively. The DAPI image was recorded to identify both the nucleus of the CTL and the image plane. In a z-stack analysis 3-10 images were recorded with a step size of 100 nm to scan the cells of interest. ZEN 2012 software (Zeiss) was used for data acquisition and image processing to achieve higher resolution.

### Immuno-staining for STED microscopy

Cells were seeded on coverslips, which were coated with poly-L-ornithine (0.1 mg/ml) and mouse anti-CD3 UCHT1 (30 µg/ml, BioLegend # 300402), for 10 min at RT (20°C ± 2°C) to allow IS formation. They were then fixed for 20 min with 4% PFA at RT and autofluorescence was quenched by 10 min incubation with 50 mM Glycine. The cells were permeabilized and blocked with 0.1% Triton X and 5% normal goat serum (NGS) in PBS for 10 min. To prevent unspecific binding of secondary antibody on the mouse anti CD3 antibodies, the samples were blocked for one hour with 5% NGS in PBS, washed and treated for an additional hour with Fab’ anti-mouse (1:50 in PBS with 5% NGS, Rockland # 8111102). Then the cells were washed and incubated with the specified primary abs (Table S1). After two washing steps, the samples were incubated with secondary abs for 45 min at room temperature (20 ± 2 °C). After several washing steps the samples were mounted with abberior mount antifade mounting medium (abberior #MM-2013-2X15ML).

### Stimulated emission depletion microscopy

Imaging was performed with a four-colour STED QUADScan (Abberior instruments GmbH) using 485 nm/0.85 mW, 561 nm/2 mW, and 640 nm/12 mW excitation pulsed lasers, and a 775 nm / 1.25 W STED laser. The pinhole size was set to 90 µm (1.13 arbitrary unit) and the probes were visualized with a 100X/NA 1.4 objective (UPLSAPO100XO, Olympus). The following acquisition protocol was applied using Imspector software (Abberior instruments GmbH). First, a single confocal section was recorded at 488 nm, 561 nm, and 640 nm to identify cells containing GzmB, TSP1-Flag, and TSP4-HA staining. Then, one section at the IS was acquired in confocal mode at 488 nm for GzmB and in 2DSTED mode at 561 nm and 647 nm to visualize the TSP1 and TSP4 staining.

The laser power was 20%, 5% and 10% at 488, 561 and 640 nm, respectively. The STED laser emitted 50% of the maximal power of 1200 mW (corresponding to 73–83 mW in the focus, with a repetition rate of 40 MHz). The pixel size was 20×20 nm, pixel dwell time was 5 µs, and the line accumulation 4. Finally, a stack of 1 µm depth was recorded of the same IS in confocal mode for GzmB and in 3DSTED mode for TSP1-Flag and TSP4-HA. The laser powers were 10, 3 and 7 % respectively. The STED laser emitted at 60% (corresponding to 87–99 mW in the focus, with a repetition rate of 40 MHz, set at 50% of 3D capacity). The voxel size was 20×20×50 nm, pixel dwell time was 5 µs, and the line accumulation 2. Laser power had to be slightly adjusted depending on the combination of secondary antibodies that were used.

### STED image analysis

Single plane images acquired at the IS, in which GzmB labelling was acquired in confocal mode and TSP-1 and TSP-4 labelling were acquired in 2DSTED mode, were segmented using Cellpose 2.0 (Pachitariu and Stringer, 2022). Images were background subtracted using ImageJ (Schindelin, 2012) then the „Fire“ look up table was applied and the image was save as RGB. We used the “cyto” model as staring point to train Cellpose 2.0 on our images over 4 to 5 rounds. Training was performed separately for each channel. After training the images were segmented automatically. In few cases manual adjustments had to be made. The segmented images were saved and subsequently used in an in house written Matlab (MathWorks) program classifying the overlapping objects according to their staining. Objects were deemed to be the same within the different channels if at least 10% of their surface overlapped.

### Tracking TSP-4 and TSP-1 transport to LGs by the retention using selective hooks (RUSH) system

3×10^6^ of 3-day activated CD8^+^ T cells were transfected using the Human T Cell Nucleofector Kit (Amaxa Biosystem) and the Amaxa Nucleofector II system (Lonza), Program T-023 with 1.5/10^6^. Cells were cultured with 50 U/ml of rhIL-2 in RPPMI-1640 medium (#8758, Merck) supplemented with 7.5% iron-enriched BCS (GE Healthcare HyClone) for 24 h and then used for a time-course of release by adding biotin (#4698, Merck) at a final concentration of 40 μM as previously described (Capitani et al., 2023). A sample of transfected cells incubated 60 min in medium without biotin was used as negative control. Cells were fixed with 4% PFA in PBS 1X (v/v) for 15 min at room temperature, permeabilized and stained with primary abs (Table S2) in 1% (w/v) BSA, 0.1 (v/v) % saponin in PBS 1X over-night at 4°C, finally incubated with fluorescent secondary abs (Table S2) for 45 min at room temperate.

### Cell culture and transfection of human CTLs for dSTORM microscopy

Isolation of peripheral blood CD8+ T cells from the buffy coat of healthy donors using the RosetteSep™ Human CD8^+^ T cell Enrichment Cocktail, as per manufacturers protocol, and subsequent centrifugation at 1,200 xg from 20 minutes with no brake at RT in a Ficoll gradient. CD8^+^ resting population are washed in R10 (RPMI-1640 medium (#7388, Merck) with 2 mM L-Glutamine, supplemented with 10 % FBS, 2 mg/ml penicillin, with 100 U/ml of rhIL-2) to remove residual Ficoll prior to culturing in R10 at 1×10^6^ cell/mL. CD8^+^ resting T cells are activated using Dynabeads™ Human T-activator CD3/CD28 in a 1:1 ratio with cells. Cells are seeded into a flat bottom 24 well plate at 1×10^6^ cell/mL including Dynabeads and incubated at 37 °C in a humidified 5% CO_2_ incubator. Beads are removed 48 hours after activation and then used for nucleofection.

Nucleofection is performed using the Amaxa T Cell nucleofector kit. 2×10^6^ cells per condition are centrifuged at 200xg for 10 minutes at RT. Resuspend in 100µL of RT Human T Cell Nucleofector Solution then combined with 1.5 µg of total DNA (TSP-1-GFPSpark and TSP-4-mCherry). The cell/DNA suspension is placed in the nucleofection cuvette and run using the T-023 Nucleofector program. Pre-equilibrated R10 culture medium is added to the cuvette then transferred into a 24 well plate to a final volume of 1mL (R10 with 50u/mL rIL2)/well/sample. Nucleofected cells are rested for 24 h, checked for double construct expression, then deposited on the planar supported lipid bilayer as previously described.

### 2D single molecule localization microscopy (SMLM) and dSTORM

The Oxford NanoImager (ONI) microscope was used to generate 2D SMLM images using dSTORM (direct Stochastic Optical Reconstruction Microscopy). A 100X objective lens/1.4 NA in oil-immersion was used. 405 nm, 488 nm, 561 nm and 640 nm lasers were used with 2 channels (with dichroic mirror split at 640 nm). All calibrations, controls and images were performed at 27°C with a super-critical TIRF angle of 54°.

After flushing out CTLs following 1.5 h of SMAP deposition on the PSLB (ICAM1 + anti-CD3ε UCHT1 Fab^’^), channels were first fixed for 30 min at 4°C with 4% PFA + 0.25% Glutaraldehyde, washed 6 times with MilliQ, then incubated with anti-RFP-AF555 and anti-GFP-AF488 overnight to optimize sample blinking during image acquisition. Ibidi channels were washed 6 times with MilliQ and then incubated with WGA-AF647 for 8 hours, then washed again. Samples were imaged in dSTORM buffer (320 µL 1x PBS, 40 µL 50% glucose, 40 µL 2-Mercaptoethylamine•HCl (2-MEA, 4 µL glucose-oxidase). First, the 640 nm laser was used at 75% laser power to excite WGA-AF647, then the 561 nm laser was used at 80% laser power to excite AlexaFluor 555, followed by the 488 nm laser to excite AlexaFluor 488. The 405 nm laser was used to promote fluorophore blinking. 3,000 frames were acquired per fluorophore. An exposure of 30 ms was used. Calibration channel mapping performed using TetraSpec Microspheres of 100 nm were used to align all three channels and generate the final three-coloured-dSTORM image using CODI software.

The Oxford NanoImager (ONI) analysis platform, CODI, was used to analyse 3 colour-dSTORM images. Localization, acquisition, and mapping files were uploaded for each image taken. Drift correction was applied using the DME algorithm. Frames were filtered first according to acquisition program settings then to exclude the initial excitation peaks for each laser. The steady state of localizations per frame was included as true blinking localizations. Filtering parameters around the sigma peak were then applied to include a sigma range of ∼50-250 nm (Gaussian fit to localize single molecules). Photon counts with a value about 300-500 were included to ensure good signal-to-noise. Localization precision filtering was then applied to include localization with precision less than or equal to 20 nm. SMAP radius and number of localizations was determined using the contained clustering algorithm using the radius and locs functions applied to the region of interest (the area of SMAP deposition at the c-SMAC). Micrographs of SMAPs were depicted with a display sigma of 5 nm using fixed representation. Files were downloaded from CODI in TIFF and PNG formats then cropped and converted to PNG micrographs for figures.

### RNA interference and CRIPSR/Cas9 genome editing

For RNAi-mediated TSP-1 and TSP-4 silencing 4×10^6^ 3-day activated CD8^+^ T cells were transfected using the Human T Cell Nucleofector Kit (Amaxa Biosystem) and the Amaxa Nucleofector II system (Lonza), Program T-023 with 150 pmol/10^6^ cells of human TSP-1 and TSP-4-specific siRNAs (#s14108 and #s14100, Thermo Fisher Scientific) and control scrambled siRNAs (#4390846) (Invitrogen™). Cells were tested for knock-down efficiency by qPCR and used for assays 24 h post-transfection. To investigate the association of TSP-1 with LGs in TSP-4 KD CTLs, we co-transfected 3×10^6^ 3-day activated CD8^+^T cells with 150 pmol/10^6^ cells of human TSP-4-specific siRNAs (#s14100, Thermo Fisher Scientific) and 1.5 μg/10^6^ of the pMax-TSP-1-GFPSpark construct. Cells were tested for TSP-1-GFPSpark expression by flow cytometry and for knock-down efficiency by qPCR 24 h post-transfection, and they were fixed and labelled for immunofluorescence microscopy.

For CRISPR-Cas9 genome editing 2×10^6^ 2-day activated CD8^+^ T cells were transfected using ribonucleoprotein complexes prepared by mixing 10 μg of Alt-R® S.p. Cas9 Nuclease V3 protein (Integrated DNA Technologies, #1081059) and 6 μg of target-specific guide RNAs (gRNA) for TSP-1 and TSP-4 (Table S1), which were designed using the web tool CRISPOR.org (Concordet&Haeussler, 2018), transcribed *in vitro* using HiScribe™ T7 High Yield RNA Synthesis Kit (#E2040S, NEB) and purified with RNA Clean & Concentrator™ (#R1017, Zymo Research). Cells were grown in complete R10 medium supplemented with 500 U/ml of rhIL-2 for 3 days, then tested for knockout efficiency by immunoblotting.

### Analysis of CTL-mediated and SMAP-mediated cytotoxicity

B cell lines used as target cell lines (i.e. Raji and MEC1) were grown at 37°C and 5% CO_2_ in RMPI-1640 medium (Merck) supplemented with 7.5% iron-enriched BCS (GE Healthcare HyClone).

For real-time calcein release-based analysis of cell-mediated cytotoxicity (Fig. 7B,C) (Kummerow et al. 2014; Chang et al., 2018), MEC1 B cells were stained with 500 nM calcein-AM (#C1430, Invitrogen™) in serum-free AIM V™ medium (#12055-091; Gibco™) with 10 mM HEPES at room temperature for 15 min, then washed twice and pulsed with a mixture of staphylococcal superantigens A, B, and E (SAgs, 2 μg/ml of each) for 1 h. 30×10^4^ MEC1 cells were plated in a 96-well black plates with clear bottom (BD Falcon) and CTLs were added at a effector-target (E:T) ratio of 3:1. Cytotoxicity were measured at 37°C and 5% CO_2_ every 10 min for 4 h using a Synergy HTX multi-mode plate reader (BioTek) at 485 nm excitation and 528 nm emission wavelengths in the bottom reader mode. Cytotoxicity (% target cell lysis) was calculated as follows: (F_live_ − γ x F_exp_) / (F_live_ − F_lyse_) x 100, where F_live_ is the fluorescence of targets alone, F_exp_ are target+CTL samples and F_lyse_ is the maximal target lysis in the presence of 1% Triton X-100 (Calbiochem). The γ value was measured at time zero: γ = F_live_ (0)/F_exp_ (0). All the experiments were performed in duplicate and averaged to obtain a value per experiment.

For the flow cytometry analysis of cell-mediated cytotoxicity (Fig. 8B; Fig. S1D) (Kabanova et al., 2016), Raji B cells were stained with 1.5 μM carboxyfluorescein diacetate succinimidyl ester (CFSE; #C34554; Invitrogen™) for 8 min at room temperature in PBS and then pulsed with a mixture of staphylococcal superantigens A, B, and E (SAgs, 2 μg/ml of each) for 1 h in serum-free AIM V™ medium (#12055-091; Gibco™). Unpulsed CFSE-stained Raji B cells were used as negative control (- SAgs). 25×10^4^ Raji B cells were mixed up with 7-day conditioned CTLs at different effector:target ratios (2.5:1, 5:1, 10:1) in 50 μl of AIM V™ medium and incubated at 37°C and 5% CO_2_ for 4 h. Thereafter, samples were diluted to 200 μl with cold PBS and acquired using a Guava Easy-Cyte flow cytometer (Merck Millipore) after adding 20 μg/ml propidium iodide (PI; #537059, Merck) to stain dead cells. Target cells alone and target cells lysed with 1% Triton X-100 (Calbiochem) were used for instrument set up and gating CFSE^+^PI^+^ (Fig. S8 for gating strategy). Cytotoxicity (% target cell lysis) was calculated as follows: (CFSE^+^PI^+^ cells − CFSE^+^PI^+^ cells in control sample) × 100 / (100 − CFSE^+^PI^+^ cells in control sample).

For the flow cytometry analysis of SMAP-mediated cytotoxicity (Fig. 7D,8D; Fig. S7B, sticky 6-channel slides (sticky-Slide VI 0.4, Ibidi) were glued to cleanroom cleaned coverslips 0.170×0.005 mm thickness (SCHOTT MINIFAB Diagnostics – NEXTERION® Glass) and coated with 10 μg/ml rhICAM-1/Fc Chimera (#720-IC-200, R&D), in presence (stimulated condition) or absence (unstimulated condition) of 5 μg/ml anti-human CD3ε (clone OKT3; #317302, BioLegend) in PBS at 37°C for 2h. Thereafter, 1-0.5-1×10^6^ CTLs/well were plated onto immobilized ICAM-1 or ICAM+anti-CD3ε mAb for 90 min at 37°C and 5% CO_2_, then CTLs were flushed out for three times with ice-cold PBS and released SMAPs, which had been captured on coated glass surfaces, were incubated with 1-0.5-1×10^6^ MEC1/well for 16 h at 37°C and 5% CO_2_. After an over-night incubation, MEC1 cells were recovered, stained with 20 μg/ml PI (Merck) and dead cells analyzed by flow cytometry. Unstained cells were used as negative control. Target cell death was calculated as the percentage of PI^+^ cells gated based on the negative control.

### Conditioning of CD8^+^ T cells by CLL cell supernatants

Culture supernatants were prepared by growing 5×10^7^ B cells purified from peripheral blood of either healthy donors or CLL patients by negative selection using RosetteSep™ Human B-cell enrichment Cocktail (#15064; STEMCELL Technologies), in 15 ml RPMI-1640 medium (#8758, Merck) supplemented with 7.5% BCS (GE Healthcare HyClone) for 48 h. Samples were centrifuged and supernatants were stored at -80°C.

Freshly isolated CD8^+^ cells were activated with Dynabeads™ Human T-Activator CD3/CD28 (Gibco™) at a cell/bead ratio of 1:0.5 for 48 h in conditioned media obtained by combining R10 medium and culture supernatants derived from either leukemic or healthy B cells (1:1 ratio). After 48 h, beads were removed and activated CD8^+^ T cells were expanded in conditioned media supplemented with fresh rhIL-2 for further 3-5 days.

### Statistics

Statistical analyses were performed using GraphPad Prism software version 6.07 (GraphPah). In figure legends N represents the number of biological replicates (donors) and n is the number of technical replicates (cells, granules and SMAPs analysed). At least three independent experiments, except for data shown in Fig. 3, 4, 5B-C, 7D-F, were performed for each assay. Normality of data distribution was tested using Anderson-Darling test, D’Agostino-Pearson omnibus normality test, Shapiro-Wilk normality test and Kolmogorov-Smirnov normality test with Dallal-Wilkinson-Lilliefor P value and normality assumption was accepted only when all datasets follow a normal distribution. Unpaired student’s t-test (unpaired) and one-way/two-way ANOVA tests followed by post-hoc Tukey correction test and multiple comparisons were used to determine the significance between two or multiple groups of datasets with normal distribution. Mann-Whitney *U* test or Kruskal-Wallis nonparametric test followed by post-hoc Dunn’s correction test were used to determine the significance between two or multiple groups of non-normally distributed datasets. One-sample t test was used to compare the mean of one or more samples to a known standard mean of 1 or 100 assigned to control samples. The number of repeats and the number of cells analyzed per sample are specified in each figure legend. Statistical analyses were performed using the Prism software (GraphPad Software). Statistical significance was defined as: ns p > 0.05; * p < 0.05; ** p < 0.01; *** p < 0.001; **** p < 0.0001.

## Acknowledgements

The authors wish to thank Meltem Hohman, Margarete Klose, Lina Chen and Stefan Balint for their valuable suggestions and for sharing protocols, Annamaria Fusillo and Elke Kurz for technical assistance, Beate Neumann for siRNA selection, and Salvatore Valitutti for critical reading of the manuscript.

## Data availability

All data generated in this study are available upon reasonable request from the corresponding author.

## References

Adams, J. C. and Lawler, J. (2011). The thrombospondins. Cold Spring Harb Perspect Biol 3, a009712.

Alawar, N., Schirra, C., Hohmann, M. et al. A solution for highly efficient electroporation of primary cytotoxic T lymphocytes. BMC Biotechnol 24, 16 (2024). 10.1186/s12896-024-00839-4

Apollonio, B., Ioannou, N., Papazoglou, D. and Ramsay, A. G. (2021). Understanding the Immune-Stroma Microenvironment in B Cell Malignancies for Effective Immunotherapy. Front Oncol 11, 626818.

Arruga, F., Gyau, B. B., Iannello, A., Vitale, N., Vaisitti, T. and Deaglio, S. (2020). Immune Response Dysfunction in Chronic Lymphocytic Leukemia: Dissecting Molecular Mechanisms and Microenvironmental Conditions. Int J Mol Sci 21.

Balint, S., Muller, S., Fischer, R., Kessler, B. M., Harkiolaki, M., Valitutti, S. and Dustin, M. L. (2020). Supramolecular attack particles are autonomous killing entities released from cytotoxic T cells. Science 368, 897–901.

Stacchini, A., M Aragno, M., Vallario, A., Alfarano, A., Circosta, P., Gottardi, D., Faldella, A., Rege-Cambrin, G., Thunberg, U., Nilsson, K. and Caligaris-Cappio, F. (1999). MEC1 and MEC2: two new cell lines derived from B-chronic lymphocytic leukaemia in prolymphocytoid transformation. Leuk Res 23, 127–36.

Blair, P. and Flaumenhaft, R. (2009). Platelet a-granules: Basic biology and clinical correlates. Blood Rev 23, 177–189.

Bolte, S. and Cordelieres, F. P. (2006). A guided tour into subcellular colocalization analysis in light microscopy. J Microsc 224, 213–32.

Boncompagni, G., Tatangelo, V., Lopresti, L., Ulivieri, C., Capitani, N., Tangredi, C., Marotta, G. Frezzato, F., Visentin, A., Ciofini, S., Gozzetti, A., Calzada-Fraile, D., Martin Cofreces, N.B., Trentin, L., Patrussi, L., Baldari, C.T. (2023). Leukemic cell-secreted interleukin-9 suppresses cytotoxic T cell-mediated killing in chronic lymphocytic leukemia. Cell Death Dis 15,144.

Boncompain, G., Divoux, S., Gareil, N., de Forges, H., Lescure, A., Latreche, L., Mercanti, V., Jollivet, F., Raposo, G. and Perez, F. (2012). Synchronization of secretory protein traffic in populations of cells. Nat Methods 9, 493–498.

Bossi, G., Stinchcombe, J. C., Page, L. J. and Griffiths, G. M. (2000). Sorting out the multiple roles of Fas ligand. Eur J Cell Biol 79, 539–43.

Bredel, M., Bredel, C., Juric, D., Harsh, G. R., Vogel, H., Recht, L. D. and Sikic, B. I. (2005). High-resolution genome-wide mapping of genetic alterations in human glial brain tumors. Cancer Res 65, 4088–96.

Capitani, N., Cassioli, C., Ravichandran, K., Baldari, C.T. (2023). Exploiting the RUSH system to study lytic granule biogenesis in CTLs. Meth Mol Biol 2654, 421–436.

Cassioli, C. and Baldari, C. T. (2022). The Expanding Arsenal of Cytotoxic T Cells. Front Immunol 13, 883010.

Chang, H. F., Schirra, C., Ninov, M., Hahn, U., Ravichandran, K., Krause, E., Becherer, U., Balint, S., Harkiolaki, M., Urlaub, H. et al. (2022). Identification of distinct cytotoxic granules as the origin of supramolecular attack particles in T lymphocytes. Nat Commun 13, 1029.

Dabir, P., Marinic, T. E., Krukovets, I. and Stenina, O. I. (2008). Aryl hydrocarbon receptor is activated by glucose and regulates the thrombospondin-1 gene promoter in endothelial cells. Circ Res 102, 1558–65.

Demetriou. P., Abu-Shah, E., Valvo, S., McCuaig, S., Mayya, V., Kvalvaag, A., Starkey, T., Korobchevskaya, K., Lee, L.Y.W., Friedrich, M., et al. (2020). A dynamic CD2-rich compartment at the outer edge of the immunological synapse boosts and integrates signals. Nat Immunol 21, 1232–43.

Frangogiannis, N. G. (2012). Matricellular proteins in cardiac adaptation and disease. Physiol Rev 92, 635–88.

Frolova, E. G., Sopko, N., Blech, L., Popovic, Z. B., Li, J., Vasanji, A., Drumm, C., Krukovets, I., Jain, M. K., Penn, M. S. et al. (2012). Thrombospondin-4 regulates fibrosis and remodeling of the myocardium in response to pressure overload. FASEB J 26, 2363–73.

Girard, F., Eichenberger, S. and Celio, M. R. (2014). Thrombospondin 4 deficiency in mouse impairs neuronal migration in the early postnatal and adult brain. Mol Cell Neurosci 61, 176–86.

Green, D. R. and Llambi, F. (2015). Cell Death Signaling. Cold Spring Harb Perspect Biol 7.

Ramsay, A.G., Johnson, A.J., Lee, A.M., Gorgun, G., Le Dieu, R., Blum, W., Byrd, J.C., Gribben, J.G. (2008). Chronic lymphocytic leukemia T cells show impaired immunological synapse formation that can be reversed with an immunomodulating drug. J Clin Invest 118, 2427–37.

Guo, N., Zabrenetzky, V. S., Chandrasekaran, L., Sipes, J. M., Lawler, J., Krutzsch, H. C. and Roberts, D. D. (1998). Differential roles of protein kinase C and pertussis toxin-sensitive G-binding proteins in modulation of melanoma cell proliferation and motility by thrombospondin 1. Cancer Res 58, 3154–62.

Henkin, J. and Volpert, O. V. (2011). Therapies using anti-angiogenic peptide mimetics of thrombospondin-1. Expert Opin Ther Targets 15, 1369–86.

Lawler, J., Derick, L. H., Connolly, J. E., Chen, J. H. and Chao, F. C. (1985). The structure of human platelet thrombospondin. J Biol Chem 260, 3762–72.

Lawler, J. and Detmar, M. (2004). Tumor progression: the effects of thrombospondin-1 and -2. Int J Biochem Cell Biol 36, 1038–45.

Lawler, J., Sunday, M., Thibert, V., Duquette, M., George, E. L., Rayburn, H. and Hynes, R. O. (1998). Thrombospondin-1 is required for normal murine pulmonary homeostasis and its absence causes pneumonia. J Clin Invest 101, 982–92.

Liang, Y., Diehn, M., Watson, N., Bollen, A. W., Aldape, K. D., Nicholas, M. K., Lamborn, K. R., Berger, M. S., Botstein, D., Brown, P. O. et al. (2005). Gene expression profiling reveals molecularly and clinically distinct subtypes of glioblastoma multiforme. Proc Natl Acad Sci U S A 102, 5814–9.

Livak, K. J. and Schmittgen, T. D. (2001). Analysis of relative gene expression data using real-time quantitative PCR and the 2(-Delta Delta C(T)) Method. Methods 25, 402–8.

Lucy, L.B. (1974). An iterative technique for the rectification of observed distributions. Astronomical Journal. 79, 745–754.

Ma, X. J., Dahiya, S., Richardson, E., Erlander, M. and Sgroi, D. C. (2009). Gene expression profiling of the tumor microenvironment during breast cancer progression. Breast Cancer Res 11, R7.

McKenzie, B. and Valitutti, S. (2023). Resisting T cell attack: tumor-cell-intrinsic defense and reparation mechanisms. Trends Cancer 9, 198–211.

Menager, M.M., Menasche, G., Romao, M., Knapnougel, P., Ho, C.H., Garfa, M., Raposo, G., Feldmann, J., Fischer, A. and de Saint Basile, G. (2007). Secretory cytotoxic granule maturation and exocytosis require the effector protein hMunc13-4. Nat Immunol 8, 257–267.

Ming, M., Schirra, C., Becherer, U., Stevens, D. R. and Rettig, J. (2015). Behavior and Properties of Mature Lytic Granules at the Immunological Synapse of Human Cytotoxic T Lymphocytes. PLoS One 10, e0135994.

Nicholas, N. S., Apollonio, B. and Ramsay, A. G. (2016). Tumor microenvironment (TME)-driven immune suppression in B cell malignancy. Biochim Biophys Acta 1863, 471–482.

Pachitariu, M., Stringer, C. Cellpose 2.0: how to train your own model. Nat Methods 19, 1634–1641 (2022). 10.1038/s41592-022-01663-4

Palao, T., Medzikovic, L., Rippe, C., Wanga, S., Al-Mardini, C., van Weert, A., de Vos, J., van der Wel, N. N., van Veen, H. A., van Bavel, E. T. et al. (2018). Thrombospondin-4 mediates cardiovascular remodelling in angiotensin II-induced hypertension. Cardiovasc Pathol 35, 12–19.

Patrussi, L., Capitani, N. and Baldari, C. T. (2021). Interleukin (IL)-9 Supports the Tumor-Promoting Environment of Chronic Lymphocytic Leukemia. Cancers (Basel) 13.

Peters, P.J., Borst, J., Oorschot, V., Fukuda, M., Krahenbuhl, O., Tschopp, J., Slot, J.W. and Geuze, H.J. (1991). Cytotoxic T lymphocyte granules are secretory lysosomes, containing both perforin and granzymes. J Exp Med 173, 1099–109.

Pfaffl, M. W. (2001). A new mathematical model for relative quantification in real-time RT-PCR. Nucleic Acids Res 29, e45.

Qabar, A., Derick, L., Lawler, J. and Dixit, V. (1995). Thrombospondin 3 is a pentameric molecule held together by interchain disulfide linkage involving two cysteine residues. J Biol Chem 270, 12725–9.

Raugi, G. J., Olerud, J. E. and Gown, A. M. (1987). Thrombospondin in early human wound tissue. J Invest Dermatol 89, 551–4.

Richardson, W,H. (1972). Bayesian-Based Iterative Method of Image Restoration. Journal of the Optical Society of America. 62, 55–59.

Sanchez-Ruiz, Y., Valitutti, S. and Dupre, L. (2011). Stepwise maturation of lytic granules during differentiation and activation of human CD8+ T lymphocytes. PLoS One 6:e27057.

Schindelin, J.; Arganda-Carreras, I. & Frise, E. et al. (2012), "Fiji: an open-source platform for biological-image analysis", Nature methods 9(7): 676–682, PMID 22743772, doi:10.1038/nmeth.2019

Schwarz, E. C., Kummerow, C., Wenning, A. S., Wagner, K., Sappok, A., Waggershauser, K., Griesemer, D., Strauss, B., Wolfs, M. J., Quintana, A. et al. (2007). Calcium dependence of T cell proliferation following focal stimulation. Eur J Immunol 37, 2723–33.

Sorlie, T., Perou, C. M., Tibshirani, R., Aas, T., Geisler, S., Johnsen, H., Hastie, T., Eisen, M. B., van de Rijn, M., Jeffrey, S. S. et al. (2001). Gene expression patterns of breast carcinomas distinguish tumor subclasses with clinical implications. Proc Natl Acad Sci U S A 98, 10869–74.

Sottile, J., Selegue, J. and Mosher, D. F. (1991). Synthesis of truncated amino-terminal trimers of thrombospondin. Biochemistry 30, 6556–62.

Stenina, O. I., Desai, S. Y., Krukovets, I., Kight, K., Janigro, D., Topol, E. J. and Plow, E. F. (2003). Thrombospondin-4 and its variants: expression and differential effects on endothelial cells. Circulation 108, 1514–9.

Stenina-Adognravi, O. (2013). Thrombospondins: old players, new games. Curr Opin Lipidol 24, 401–9.

Stenina-Adognravi, O. (2014). Invoking the power of thrombospondins: regulation of thrombospondins expression. Matrix Biol 37, 69–82.

Sun, L., Hui, A. M., Su, Q., Vortmeyer, A., Kotliarov, Y., Pastorino, S., Passaniti, A., Menon, J., Walling, J., Bailey, R. et al. (2006). Neuronal and glioma-derived stem cell factor induces angiogenesis within the brain. Cancer Cell 9, 287–300.

Turashvili, G., Bouchal, J., Ehrmann, J., Fridman, E., Skarda, J. and Kolar, Z. (2007). Novel immunohistochemical markers for the differentiation of lobular and ductal invasive breast carcinomas. Biomed Pap Med Fac Univ Palacky Olomouc Czech Repub 151, 59–64.

