## Supplemental figures for "Thrombospondins 1 and 4 undergo coordinated transport to multicore cytotoxic granules to regulate SMAP biogenesis and function in CTL-mediated cytotoxicity"

### Supplementary figure legends and videos

**Figure S1. Generation of effector CD8$^+$ T cells.** (**A**) Timeline of the protocol used for *in vitro* generation of cytotoxic CD8$^+$ T cells acquiring cytolytic activity (day 5 and 7 after purification) from primary CD8$^+$ T lymphocytes (day 0) purified from peripheral blood of healthy donors. (**B**) *Left*, Representative dot plot showing the percentage (%) of naïve ($T_{naive}$, CD62L$^+$/CD45RA$^+$), central memory ($T_{CM}$, CD62L$^+$/CD45RA$^-$), effector memory ($T_{EM}$, CD62L$^-$/CD45RA$^-$) and terminal effector memory re-expressing CD45RA ($T_{EMRA}$, CD62L$^-$/CD45RA$^+$) in differentiated 5-day CTLs generated as depicted in (**A**). *Right*, The graph (mean±SD) shows the percentage of naïve, central memory, effector memory and late effector memory re-expressing CD45RA detected in 5-day CTL cultures of 5 different donors. (**C**) RT-qPCR of granzyme B (GZMB) and perforin (PRF) in freshly isolated CD8$^+$ T cells (day 0) and 5-day and 7-day CTLs. The graphs (mean±SD, ctr value = 1) show the abundance of the analysed transcripts, which was determined using the $\Delta\Delta$Ct method and normalized to 18S ribosomal RNA. $N_{donors}$ = 6, one sample t-test; * $p \leq 0.05$, only significant differences are shown. Each dot represents one donor. (**D**) Fluorimetric analysis of cytotoxicity mediated by CTLs using a calcein release-based assay. The curve shows the kinetics of target cell death by CTLs generated as depicted in A at the effector:target ratio of 3:1. $N_{donors}$ = 2. (**E,F**) Validation of anti-TSP-4/1 antibodies in KO CTLs. Immunoblot analysis of TSP-4 (E) and TSP-1 (F) in lysates of control and TSP-4 KO or TSP-1 KO CTLs gene-edited by CRISPR-Cas9 technology. Actin was used as loading control. The migration of molecular mass markers is indicated (kDa).

**Figure S2. Analysis of TSP-1-GFPSpark, TSP-4-mCherry and TSP-4-GFPSpark expression and localization in transfected CTLs.** (**A**) Schemes of the expression constructs with 3' fluorescent tags encoding full-length TSP-1 and TSP-4 fused to the N-terminus of GFPSpark (GFPS) and mCherry (mC), respectively. (**B**) Representative FACS profile of CTLs expressing mC alone or TSP-4-mC and stained with an anti-RFP antibody. (**C**) Representative FACS profiles of CTLs expressing GFPS alone, TSP-1-GFPS (*left*) or TSP-4-GFPS (*right*) and stained with anti-GFP antibody. (**D**) Representative immunoblots of lysates of CTLs expressing either mC alone or TSP-4-mC. Samples were probed with anti-RFP (not shown) and anti-TSP-4 antibodies. Actin was used as a loading control. The migration of molecular mass markers is indicated (kDa). (**E**) Representative immunoblots of lysates of CTLs expressing either GFPS alone, TSP-1-GFPS (*left*) or TSP-4-GFPS (*right*). Samples were probed with anti-GFP (not shown), anti-TSP-1 and anti-TSP-4 antibodies. Actin was used as a loading control. The migration of molecular mass markers is indicated (kDa). (**F**) Confocal images (medial optical sections) of 4-day CTLs expressing either mC alone or TSP-4-mC. Dashed lines mark the cell outline. Scale bar: 5 µm. (**G**) Confocal images (medial optical sections) of CTLs expressing either GFPS alone, TSP-1-GFPS or TSP-4-GFPS. Dashed lines mark the cell outline. Scale bar: 5 µm.

**Figure S3. Human CTLs contain typical MCGs and SCG.** (**A**) Workflow for the four color post-embedding CLEM procedure**.** (**B**) Representative TEM images of six human CTLs without electroporation. Shown are TEM images of human CTLs (left). The white rectangles mark the magnified area in the images on the right. White arrows indicate typical multicore granules (MCGs) characterized by a typical dense core (SMAP). The black arrow indicates a typical single core granule (SCG) (right panel, lower image). $N_{donors}$ = 3, $n_{cells}$ = 21.

**Figure S4. GzmB-mCherry signal largely overlaps anti-GzmB staining and co-localizes with TSP-4 and TSP-1 at the IS.** (**A**) *Left*, Confocal images (medial optical sections) of CTLs expressing GzmB-mCherry and co-stained with an anti-GzmB antibody. Dashed lines mark the cell outline. Scale bar: 5 µm. *Right*, Quantifications (mean±SD) of the weighted colocalization using the Manders' overlap coefficient between GzmB-mCherry signal with anti-GzmB staining. (**B,C**) Representative maximum intensity z-projection and orthogonal views from confocal z-stacks of CTLs expressing either TSP-4-GFPSpark (green) (B) or TSP-1-GFPSpark (green) (C) and interacting with activating [ICAM1-AF405 + anti-CD3ε UCHT1 Fab' (unlabelled)] ligands (*right*) for 30 min, followed by staining with an anti-GzmB antibody (magenta). The formation of a mature IS is indicated by the presence of an ICAM-1 ring (blue). Scale bar: 5 µm. 3D reconstructions of representative z-stacks are shown in Supplementary Videos 3-4.

**Figure S5**. **Quality control and analysis pipeline of 2D SMLM dSTORM acquisitions.** (**A**) Light program settings for 2D SMLM dSTORM imaging using the ONI. The Rainbow LUT is used to depict the sequence of localisation acquisitions from 0-3000 (647), 3001-6000 (555) and 6001-9000 (488), indicated as the frame index. (**B**) The image analysis pipeline following upload of raw files into the CODI analysis platform. (**C,E**) Frame index micrographs of triple and double positive particles demonstrating localisations of each color were acquired during the appropriate laser excitation frame period (shown by the dotted lines). (**D,F**) The localization precision and sigma peak corresponding to the images of the particle in (C,E). The base of the localization precision peak at 20 nm indicates a resolution of 20 nm was achieved. The sharp, single Gaussian peak on the sigma plot indicates the image acquisition had good fluorophore blinking quality and therefore accurate molecule localization.

**Figure S6. 2D SMLM dSTORM of CTL-derived SMAPs at the IS.** (**A**) Micrographs of SMAPs deposited at the IS by CTLs on the PSLB with particles of interest denoted by grey boxes. Scale bar: 2 µm. (**B**) Table summarizing the analysis of each particle category based on average diameter, particle number and number of size outliers. (**C**) Additional representative micrographs of all three SMAP categories. SMAPs ware stained with WGA-AF647, TSP4-mCherry was probed with anti-RFP-AF555 and TSP1-GFPSpark was probed with anti-GFP-AF488. Scale bars are indicated in each panel row.

**Figure S7. TSP-4 depletion by CRISPR/Cas9 gene editing impairs SMAP-mediated cytotoxicity.** (**A**) Immunoblot analysis of TSP-4 in control (ctr) and TSP-4 KO cells. Data are expressed (mean±SD) as residual % of mRNA in KD samples compared to control. $N_{donors}$ = 5, one-sample t test; ** $p \leq 0.01$. (**B**) Flow cytometric analysis (mean±SD) of cytotoxicity mediated by the synaptic output of control (ctr) and TSP-4 KO plated on immobilized ICAM-1 and ICAM-1 + anti-CD3ε mAb. MEC1 B cells, used as targets, were seeded after flushing out the CTLs and incubated for 16 h at 37°C. Targets were then recovered and analyzed by flow cytometry using propidium iodide (PI) to stain dead cells. Quantification (mean±SD) of the target cell lysis (%) (left) and SMAP-mediated cytotoxicity expressed as fold change in KO samples versus ctr. $N_{donors} = 5$, one-way ANOVA test (left) and one-sample t test (right); ** $p \leq 0.01$, * $p \leq 0.05$, only significant differences are shown.

**Figure S8. Gating strategy used for the flow cytometric analysis of cell-mediated cytotoxicity.** (**A**) Raji B cells used as target cells were labeled with CFSE and the analysis was carried out gating on CFSE-positive ($CFSE^+$) cells. $CFSE^+$ Raji cells were further gated and the analysis of $PI^+$ cells restricted to $CFSE^+$ cells.

**Video 1-2.** The movies show representative confocal z-stacks of CTLs co-expressing TSP-4-mCherry andTSP-1-GFPSpark, plated on non-activating (**1**) and activating (**2**) PSLBs and co-stained for GzmB. ICAM-1, blue; TSP-1-GFPSpark, green; TSP-4-GFPSpark, yellow; GzmB, magenta.

**Video 3-4.** The movies show representative confocal z-stacks of CTLs co-expressing TSP-4-GFPSpark and GzmB-mCherry (**3**) or TSP-1-GFPSpark and GzmB-mCherry (**2**), plated on activating PSLBs. ICAM-1, blue; TSP-4-GFPSpark or TSP-1-GFPSpark, green; GzmB, magenta.

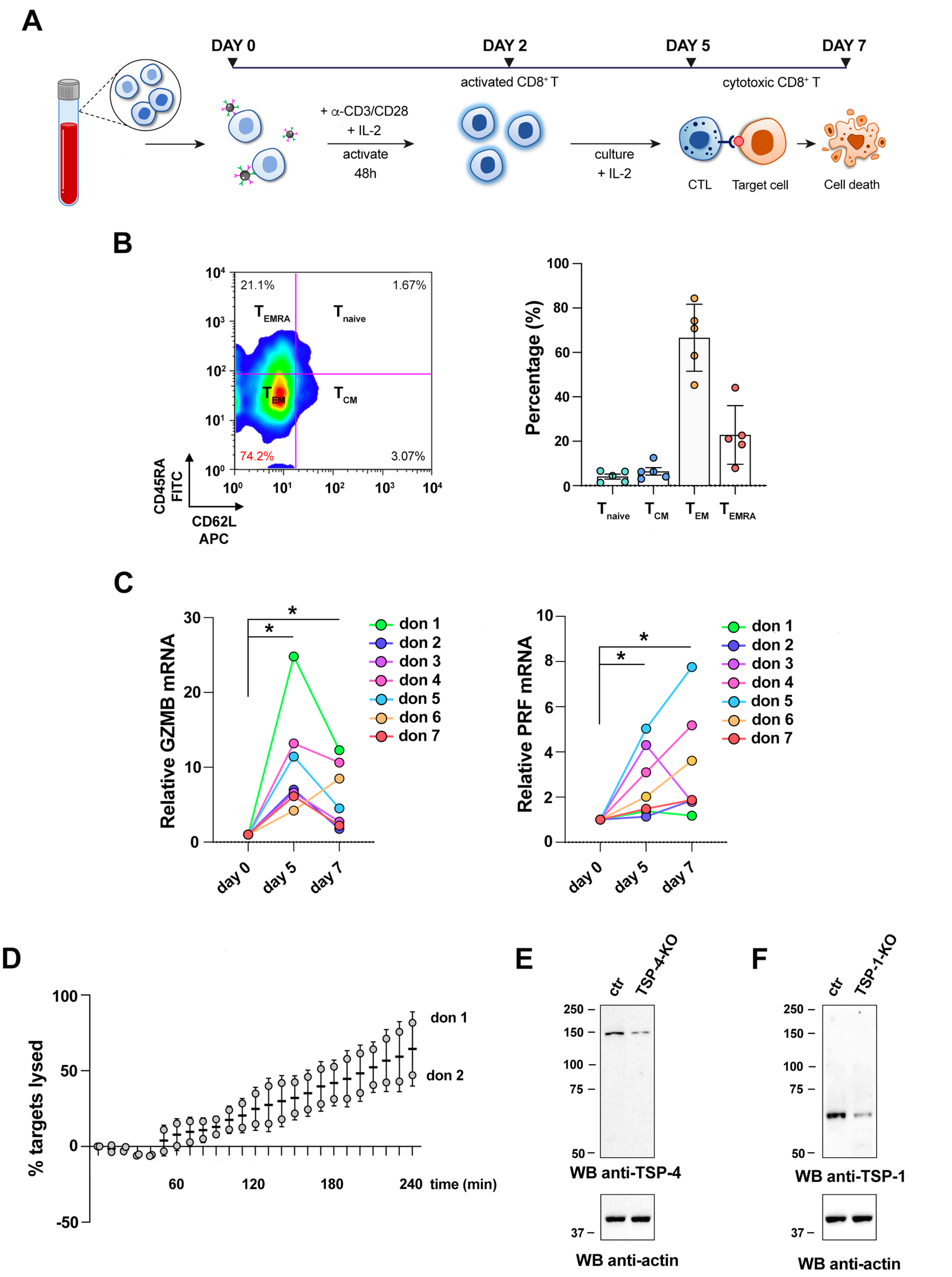
A
DAY 0
DAY 2
DAY 5
DAY 7
activated CD8+ T
cytotoxic CD8+ T
+ α-CD3/CD28
+ IL-2
activate
48h
culture
+ IL-2
CTL
Target cell
Cell death
B
21.1%
1.67%
TEMRA
Tnaive
TEM
TCM
74.2%
3.07%
CD45RA
FITC
CD62L
APC
Percentage (%)
Tnaive
TCM
TEM
TEMRA
C
Relative GZMB mRNA
Relative PRF mRNA
day 0
day 5
day 7
don 1
don 2
don 3
don 4
don 5
don 6
don 7
D
% targets lysed
time (min)
don 1
don 2
E
ctr
TSP-4-KO
WB anti-TSP-4
WB anti-actin
F
ctr
TSP-1-KO
WB anti-TSP-1
WB anti-actin

FIGURE S1

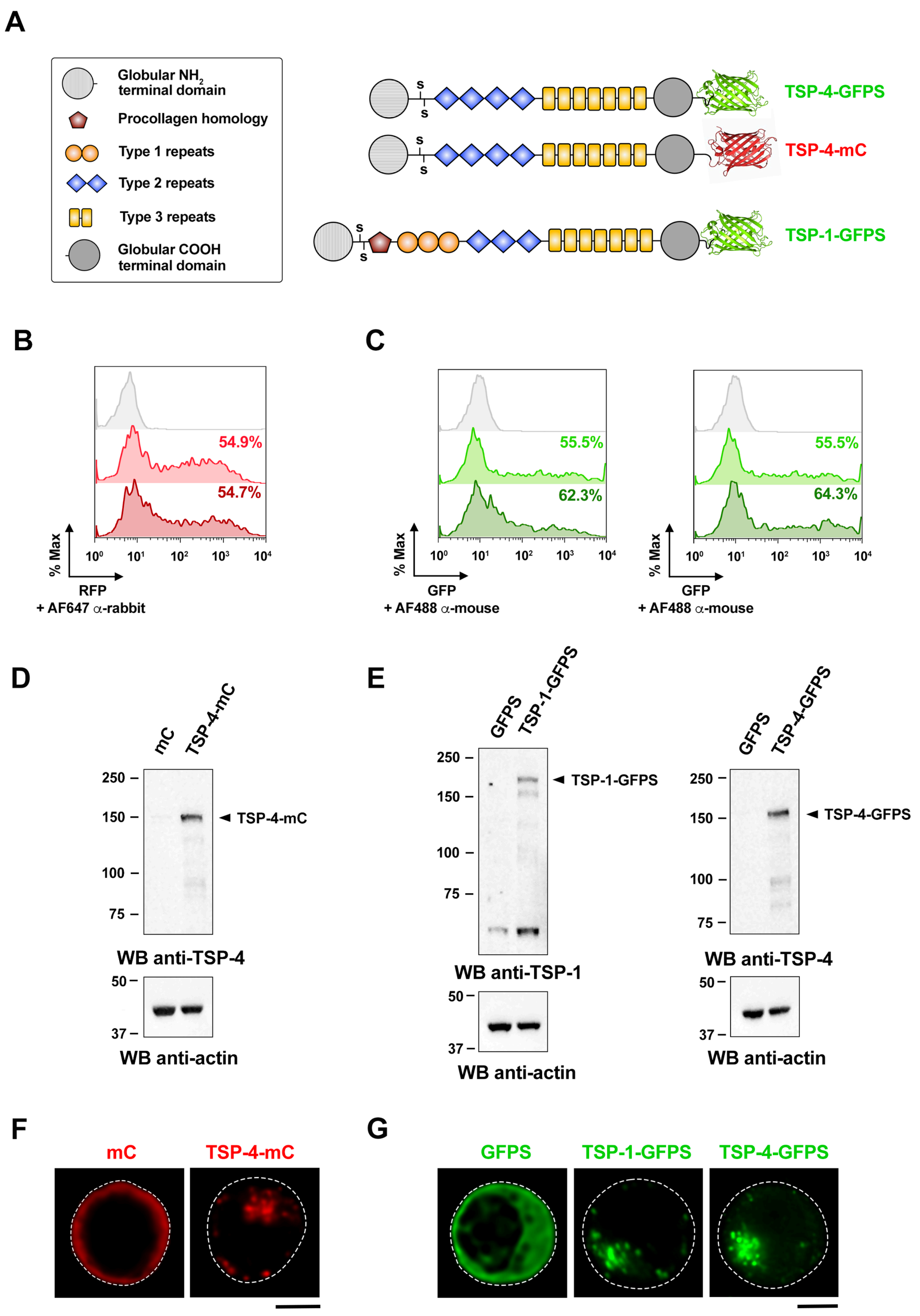
A
Globular NH2 terminal domain
Procollagen homology
Type 1 repeats
Type 2 repeats
Type 3 repeats
Globular COOH terminal domain
TSP-4-GFPS
TSP-4-mC
TSP-1-GFPS
B
54.9%
54.7%
% Max
RFP
+ AF647 α-rabbit
C
55.5%
62.3%
55.5%
64.3%
GFP
+ AF488 α-mouse
D
mC
TSP-4-mC
250
150
100
75
TSP-4-mC
WB anti-TSP-4
50
37
WB anti-actin
E
GFPS
TSP-1-GFPS
TSP-1-GFPS
WB anti-TSP-1
WB anti-actin
GFPS
TSP-4-GFPS
TSP-4-GFPS
WB anti-TSP-4
WB anti-actin
F
mC
TSP-4-mC
G
GFPS
TSP-1-GFPS
TSP-4-GFPS

FIGURE S2

**A**

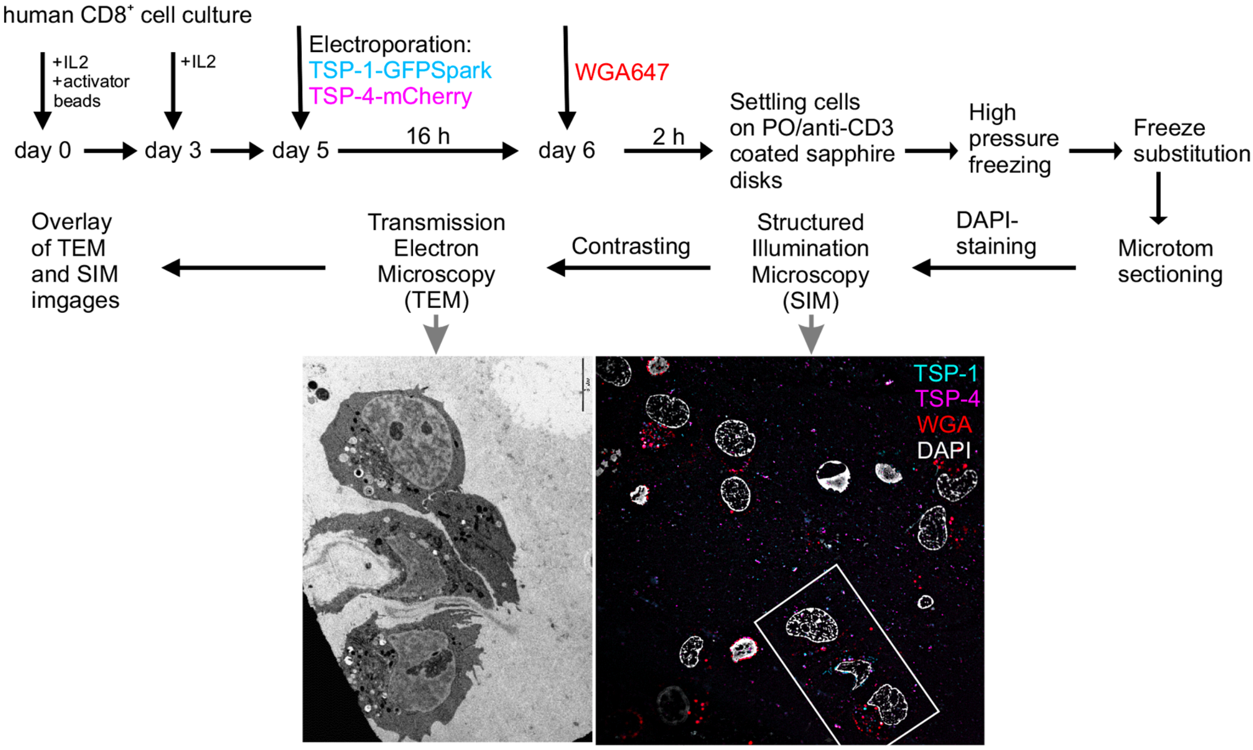

human CD8+ cell culture
+IL2
+activator beads
+IL2
Electroporation:
TSP-1-GFPSpark
TSP-4-mCherry
WGA647
day 0
day 3
day 5
16 h
day 6
2 h
Settling cells on PO/anti-CD3 coated sapphire disks
High pressure freezing
Freeze substitution
Microtom sectioning
DAPI-staining
Structured Illumination Microscopy (SIM)
Contrasting
Transmission Electron Microscopy (TEM)
Overlay of TEM and SIM imgages
TSP-1
TSP-4
WGA
DAPI


**B**

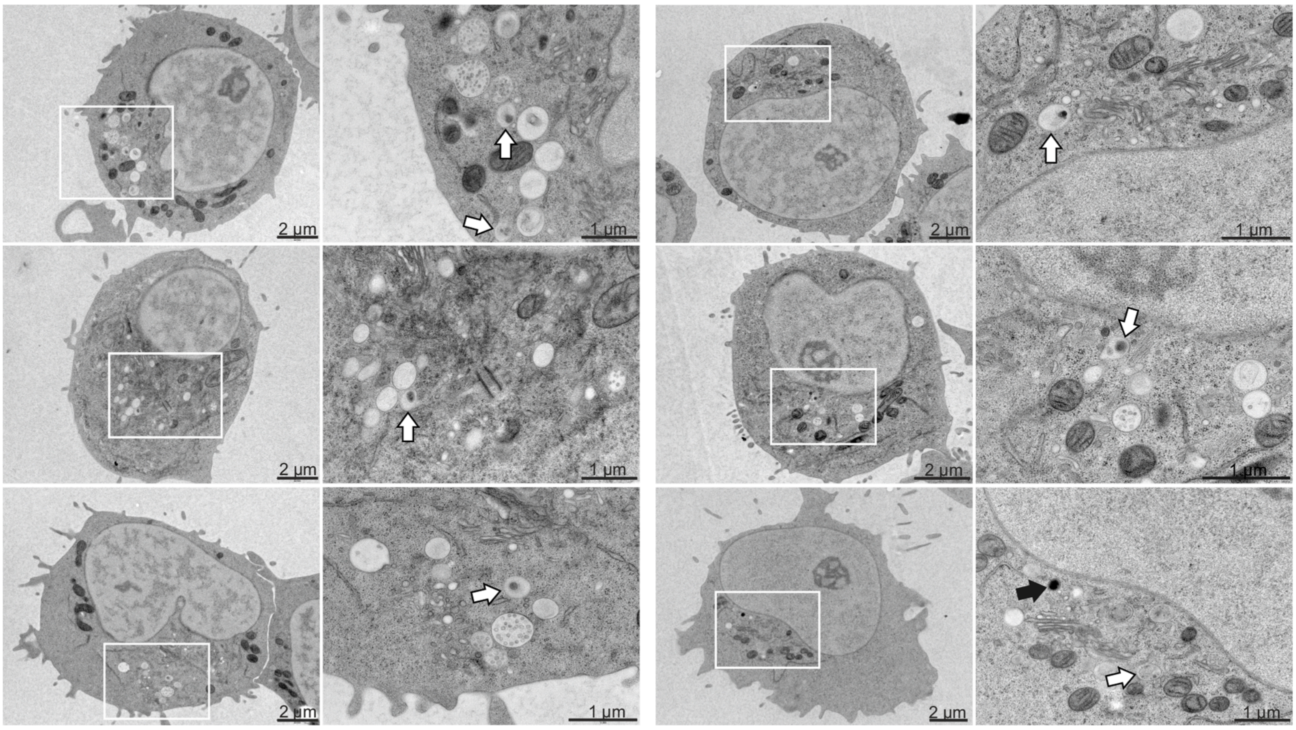

2 µm
1 µm
2 µm
1 µm
2 µm
1 µm
2 µm
1 µm
2 µm
1 µm
2 µm
1 µm


**FIGURE S3**

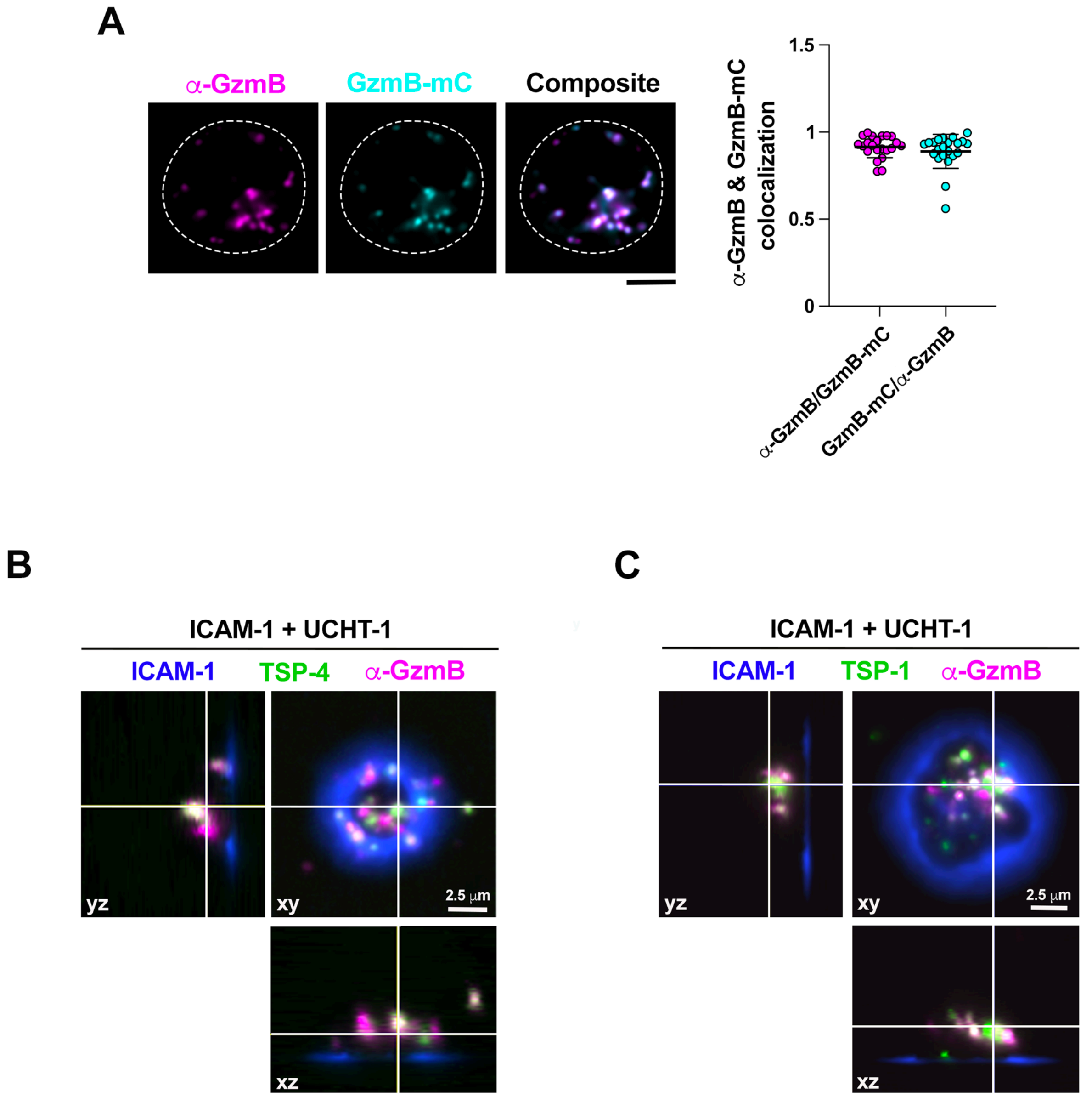
A
α-GzmB
GzmB-mC
Composite
1.5
1
0.5
0
α-GzmB & GzmB-mC colocalization
α-GzmB/GzmB-mC
GzmB-mC/α-GzmB
B
ICAM-1 + UCHT-1
ICAM-1
TSP-4
α-GzmB
yz
xy
2.5 µm
xz
C
ICAM-1 + UCHT-1
ICAM-1
TSP-1
α-GzmB
yz
xy
2.5 µm
xz

FIGURE S4

**A**

| | Light program | | |
|---|---|---|---|
| Lazer | Channel 1 - far red | Channel 0 - red/orange | Channel 2 - blue/green |
| Fluorophore | WGA-Alexafluor 647 | Tsp4-Alexafluor 555 | Tsp1-Alexafluor 488 |
| Power | 75% lazer power | 80% lazer power | 80% lazer power |
| Frames | 3000 | 3000 | 3000 |

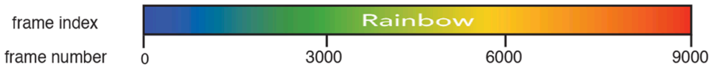


**B**

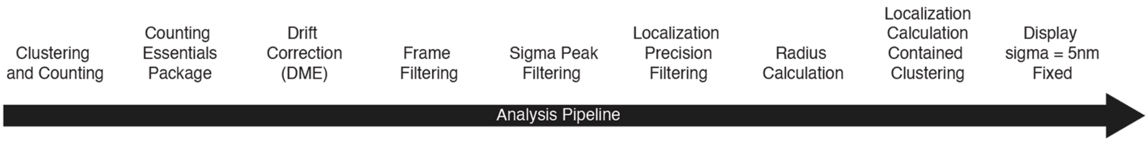


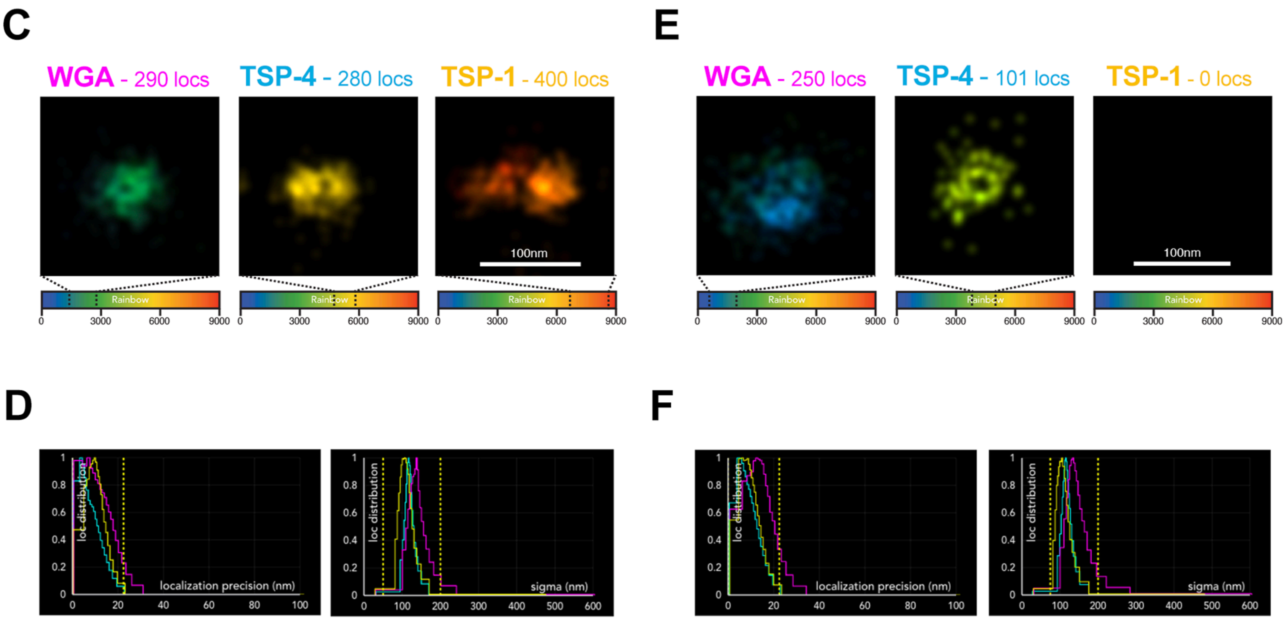


**FIGURE S5**

**A**

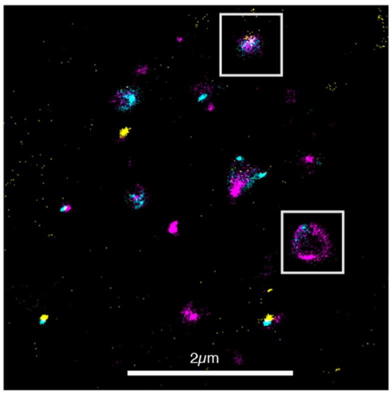


**B**

| | WGA$^+$ TSP4$^-$ TSP1$^+$ | WGA$^+$ TSP4$^+$ TSP1$^-$ | WGA$^+$ TSP4$^+$ TSP1$^+$ |
|---|---|---|---|
| Average Diameter | 140nm | 181nm | 174nm |
| Number of Particles | 9 | 16 | 45 |
| Number of Size Outliers | 0 | 0 | 1 |

**C**

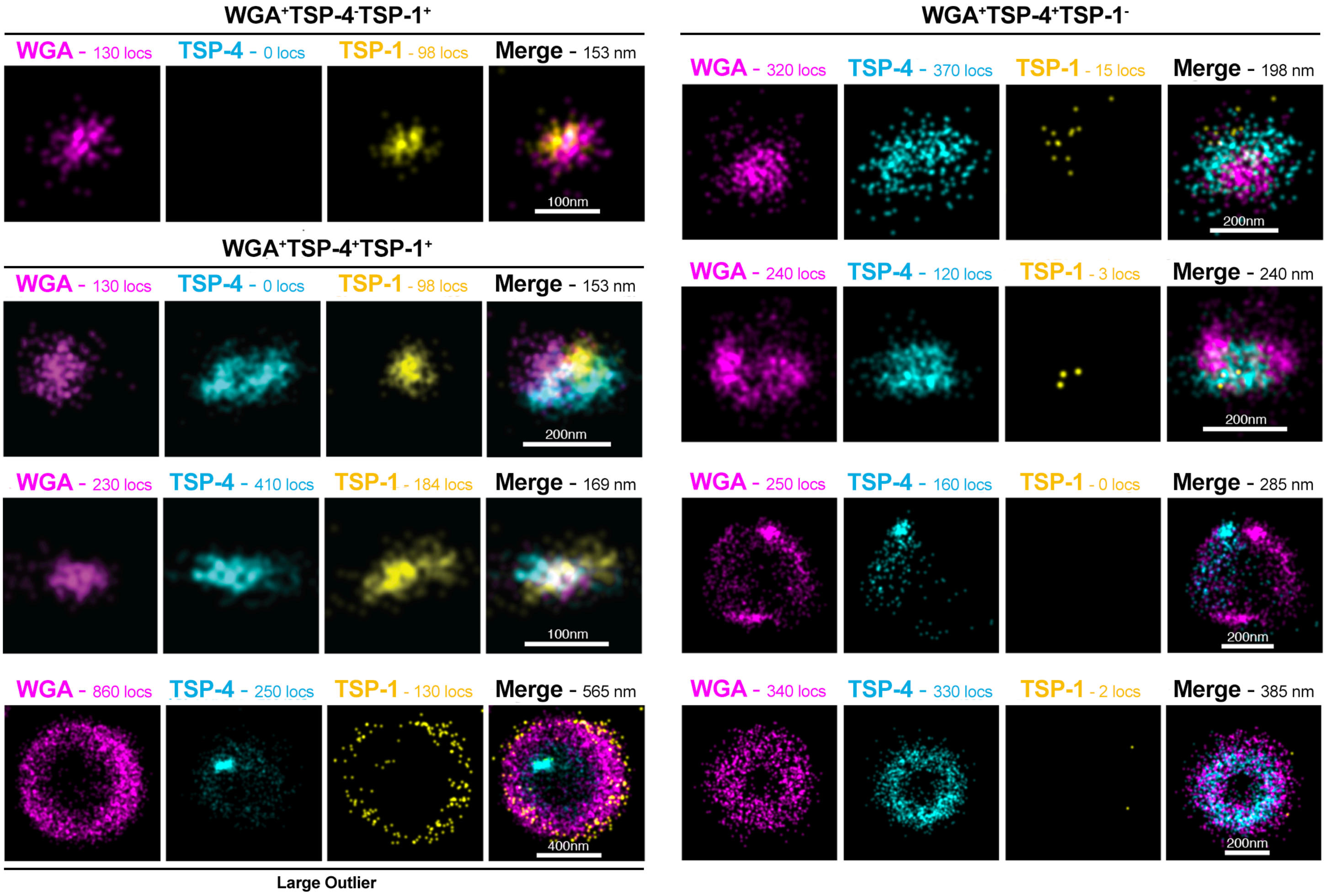


**FIGURE S6**

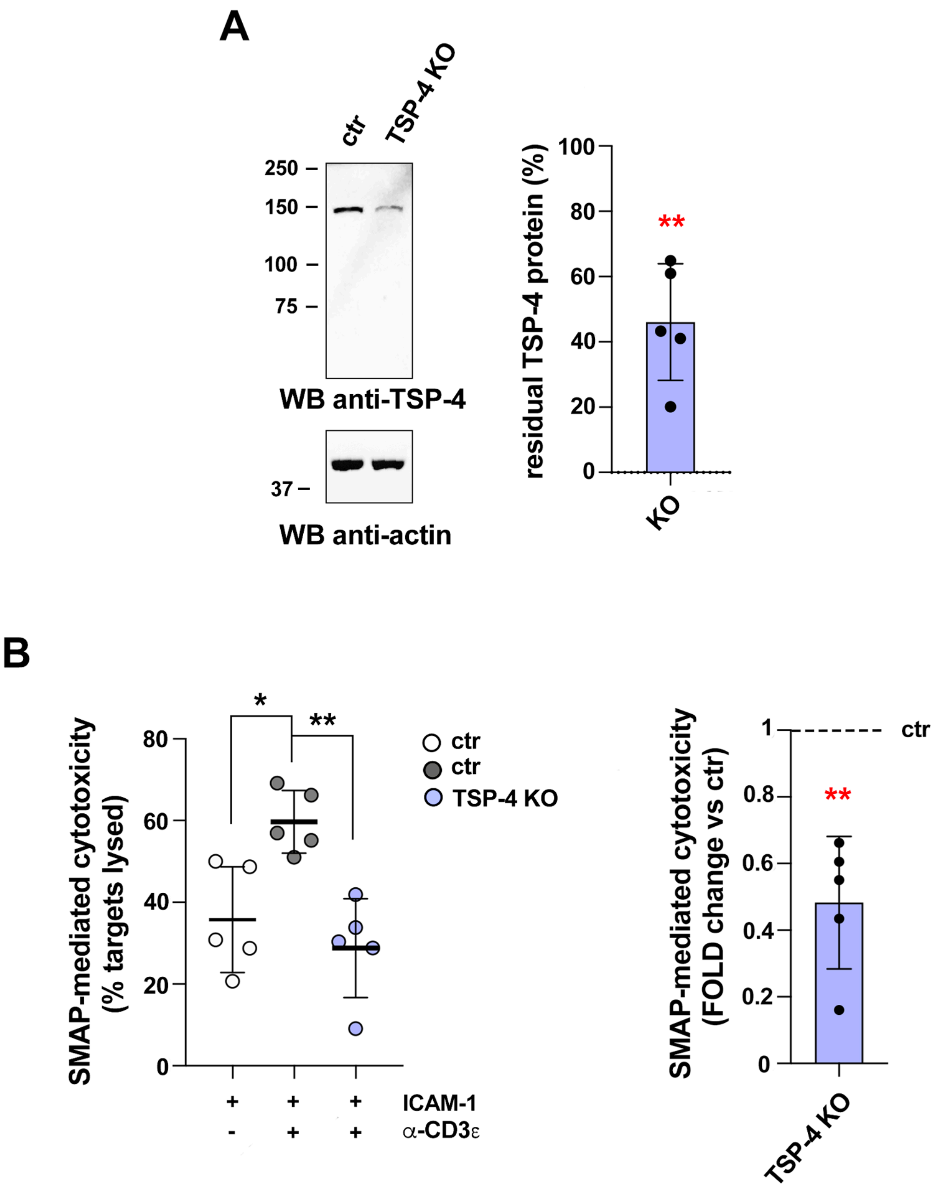
A
ctr
TSP-4 KO
250
150
100
75
WB anti-TSP-4
37
WB anti-actin
residual TSP-4 protein (%)
100
80
60
40
20
0
**
KO
B
SMAP-mediated cytotoxicity
(% targets lysed)
80
60
40
20
0
*
**
ctr
ctr
TSP-4 KO
ICAM-1
α-CD3ε
+ + +
- + +
SMAP-mediated cytotoxicity
(FOLD change vs ctr)
1
0.8
0.6
0.4
0.2
0
ctr
**
TSP-4 KO

FIGURE S7

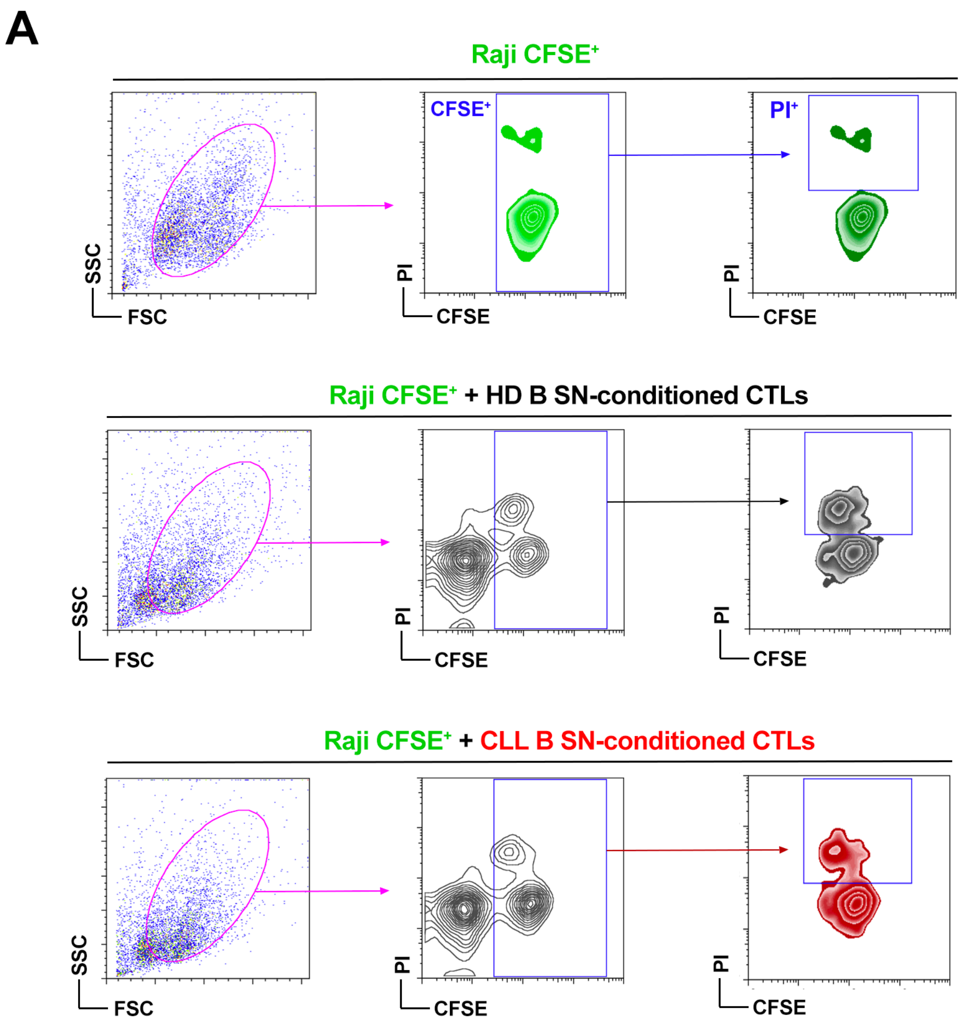
A
Raji CFSE+
CFSE+
PI+
SSC
FSC
PI
CFSE
Raji CFSE+ + HD B SN-conditioned CTLs
Raji CFSE+ + CLL B SN-conditioned CTLs

**FIGURE S8**
